# *Pseudomonas aeruginosa* contracts mucus to form biofilms in tissue-engineered human airways

**DOI:** 10.1101/2022.05.26.493615

**Authors:** Tamara Rossy, Tania Distler, Joern Pezoldt, Jaemin Kim, Lorenzo Talà, Nikolaos Bouklas, Bart Deplancke, Alexandre Persat

## Abstract

The opportunistic pathogen *Pseudomonas aeruginosa* causes antibiotic-recalcitrant pneumonia by forming biofilms in the respiratory tract. Despite extensive *in vitro* experimentation, how *P. aeruginosa* forms biofilms at the airway mucosa is unresolved. To investigate the process of biofilm formation in realistic conditions, we developed AirGels: 3D, optically-accessible tissue-engineered human lung models that emulate the airway mucosal environment. AirGels recapitulate important factors that mediate host-pathogen interactions including mucus secretion, flow and air-liquid interface, while accommodating high resolution live microscopy. With AirGels, we investigated the contributions of mucus to *P. aeruginosa* biofilm biogenesis in *in vivo*-like conditions. We found that *P. aeruginosa* forms mucus-associated biofilms within hours by contracting luminal mucus early during colonization. Mucus contractions facilitates aggregation, thereby nucleating biofilms. We show that *P. aeruginosa* actively contracts mucus using retractile filaments called type IV pili. Our results therefore suggest that, while protecting epithelia, mucus constitutes a breeding ground for biofilms.

## INTRODUCTION

Bacteria predominantly colonize their environments in the form of biofilms, dense communities of contiguous cells embedded in a self-secreted polymeric matrix^1^. The mechanisms of biofilm formation have been extensively studied on abiotic surfaces and in laboratory conditions^2,3^. In contrast, our understanding of biofilm morphogenesis in a realistic context of human infections is limited^4,5^. Biofilms from the pathogen *P. aeruginosa* epitomize this disparity. Clinical observations show that *P. aeruginosa* forms airway-associated biofilms during acute and chronic pneumoniae in immunocompromised individuals^6,7^. Due to their clinical prevalence, *P. aeruginosa* biofilms have been meticulously investigated *in vitro*. Still, the architecture of *in vivo P. aeruginosa* biofilms sampled from patient sputum and lung explants is quite distinct from *in vitro* ones^5^. This discrepancy indicates that biofilm studies in axenic environments omit critical factors of the airway mucosal surface that contribute to biofilm morphogenesis.

Epithelial tissues are lined with a hydrogel substance called mucus (Figure 1a), the first line of defense of the airway against respiratory pathogens. Dedicated goblet cells secrete gel-forming mucin glycoproteins which crosslink into a viscoelastic substance upon exocytosis to form mucus^8,9^. The mucus hydrogel mesh is impermeable to large particles, thereby functioning as a passive physical barrier^8,9^. Individuals with underlying respiratory conditions such as chronic obstructive pulmonary disease (COPD) and cystic fibrosis (CF) have aberrant mucus. At the same time, they are at risk of specifically developing chronic *P. aeruginosa* pneumonia^10^. Despite this common association, how mucus mechanics contribute to the onset and persistence of *P. aeruginosa* during infection remains unresolved.

**Figure 1:**
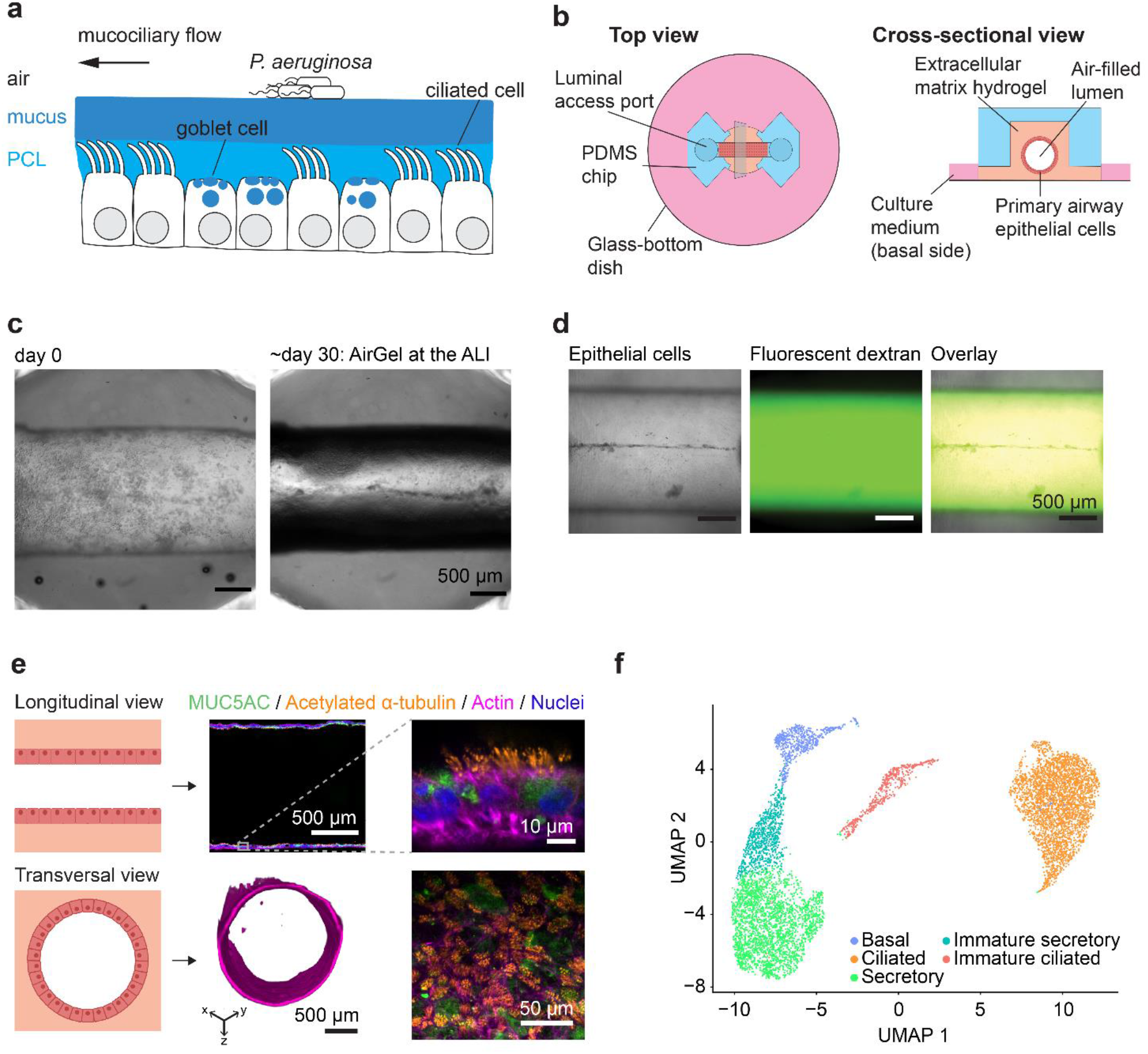
A tissue-engineered airway as a novel infection model. **(a)** Simplified representation of the airway mucosa. Mucus-secreting goblet cells and beating ciliated cells are essential to generate mucociliarly clearance, which transports inhaled pathogens out of the airway. PCL: periciliary layer. **(b)** Schematic of an AirGel chip. **(c)** Brightfield image of an AirGel on the day of HBE cell were seeded (left) and at the air-liquid interface after 30 days in culture (right). **(d)** AirGels permeability measurement by dextran assay. The brightfield image (left) shows the epithelial cells lining the lumen; the epifluorescence image (center) shows signal from the fluorescent 4 kDa dextran which does not cross the epithelial barrier, as shown in the overlay picture (right). **(e)** Longitudinal cross-sectional images of immunostained differentiated AirGels. Confocal images show the gel-forming mucin MUC5AC (green) and acetylated α-tubulin labeling cilia (orange) along with the actin dye phalloidin (pink) and nuclear dye DAPI (blue). The transverse cross section 3D image was reconstituted from SPIM data for actin fluorescence. The bottom right panel is a maximal intensity projection of a z-stack acquired in the curved lumen. **(f)** scRNA-seq identifies cell type diversity of AirGels. Uniform Manifold Approximation and Projection (UMAP) embedding of cells pooled from three differentiated AirGels, subjected to scRNA-seq profiling.

*In vitro* experimentations show that mucins influence *P. aeruginosa*’s multicellular lifestyle. Mucin-coated surfaces and concentrated mucus repress *P. aeruginosa* motility^11–13^, thereby favoring biofilm formation. Also, mucin polymers generate entropic forces that passively promote aggregation^14^. In other conditions, natively purified mucins can also have a negative effect on biofilms biogenesis by stimulating motility and dispersal^15–17^. These experiments each capture different chemical and physical aspects of mucus, but how all these contributions balance *in vivo* to influence biofilm formation has yet to be resolved. Minimally invasive experimental models that replicate physicochemical properties of the airway mucosa have the potential to bring a new perspective on this process.

Existing airway infection models have limitations that prevent mechanistic investigations of bacterial infections at the single cell level. Tracheal explants from animal models allow dynamic studies^18–23^. They are however short-lived, displaying a rapid depletion in goblet cells after a few hours with signs of apoptosis^24^. In addition, murine and human airway mucus show distinct composition and distribution^18^. 2D *in vitro* models systems based on porous membranes inserts^25–27^ and lung-on-a-chip devices^28–30^ are not suited for high-resolution microscopy and lack morphological accuracy. Organoids have a strong potential in recapitulating physical and biological aspects of the mucosal environment^31–33^. However, the cystic morphology of organoids prevents the establishment of an air-liquid interface (ALI) necessary to reproduce *in vivo* conditions. In addition, infecting organoids requires microinjection of bacterial suspension, an invasive and tedious process.

To bridge the gap between *in vitro* biofilm studies and clinical observations, we used a tissue-engineering approach to faithfully emulate the mucosal environment of the airway in the lab. We engineered AirGels (<u>air</u>way in <u>gels</u>): human lung epithelial tissues supported by a tubular collagen/Matrigel extracellular matrix (ECM) scaffold^34,35^. We demonstrate that AirGels recapitulate key features of the human airway epithelium, including accurate cell types, mucus secretion and ciliary beating. We can non-invasively infect AirGels with *P. aeruginosa* while maintaining the ALI to image biofilm formation at high spatiotemporal resolution. Using this new infection model, we found that *P. aeruginosa* forms biofilms on mucus via a previously unknown mechanism. By tracking live biofilms *in situ*, we found that *P. aeruginosa* aggregate with one another by actively contracting mucus. Using a combination of simulations and biophysical experiments in selected mutants, we show that *P. aeruginosa* uses long and thin motorized filament called type IV pili (T4P) to generate the force necessary to contract mucus.

## RESULTS

### AirGel: a tissue-engineered airway infection model

We grew AirGels from primary human bronchial epithelial (HBE) cells, which expand to confluence on the cylindrical cavity of the ECM scaffold (Figure 1b). An elastomeric microfluidic chip maintains AirGels and allows for luminal access. The ECM geometry guides epithelial architecture, enabling morphological customization of AirGels. Here, we designed and optimized AirGels to enable high resolution fluorescence microcopy to monitor infection dynamics at the single bacterium level in live tissue. Maintaining an air-liquid interface in the lumen promotes epithelial cell differentiation and reproduces the physiological conditions encountered in the airway. We therefore optimized the matrix formulation so that AirGels remain stable at the air-liquid interface, thereby biologically and physically replicating the airway environment (Figure 1c).

AirGel epithelia are tight and impermeable (Figure 1d). Single-plane illumination microscopy images show that mature AirGels form tubular epithelial tissue, recapitulating the architecture and dimensions of a human small bronchus (Figure 1e)^36,37^. We characterized HBE cell differentiation in 34-day-old AirGels. Immunofluorescence highlighted an abundant population of mucus-producing goblet cells and ciliated cells (Figure 1e). To quantify the abundance of each cell type, we performed single-cell RNA sequencing (scRNA-seq) of mature AirGels. We identified five main clusters (Figure 1f and Supplementary Figure 1): basal cells (8%), ciliated cells (41%), secretory cells (34%), as well as immature ciliated (7%) and immature secretory cells (which are also sometimes defined as suprabasal cells) (10%). AirGels therefore reproduce the cellular composition and histological signature of human airway epithelia^38–41^, and more specifically the distal human airway^42^.

Given its prominent function in host-microbe interactions, we carefully characterized the architecture of mucus in AirGels. Immunofluorescence against the airway gel-forming mucins MUC5AC and MUC5B showed the presence of extracellular mucus in the form of thick luminal filaments (Figure 2a). We also observed similar fiber-like mucus architecture in live AirGels by staining with the fluorescently-labeled lectin jacalin^21^. These fibers recapitulate the mucus architecture observed in porcine and murine tracheal explants^18,20,21^. We then characterized AirGel mucociliary clearance functions. Measurements of cilia beating frequency in AirGels were indistinguishable from previous *ex vivo* measurements (Figure 2b & Supplementary Video 1)^9,43–45^. In addition, AirGel cilia generated a directional flow whose clearance velocity matched the physiological range (Figure 2c)^9,18,23^. Overall, AirGels reproduce biological, physical and dynamic parameters of the human airway including its tube-shape, all in a system allowing for live imaging of host-pathogen interactions at high resolution.

**Figure 2:**
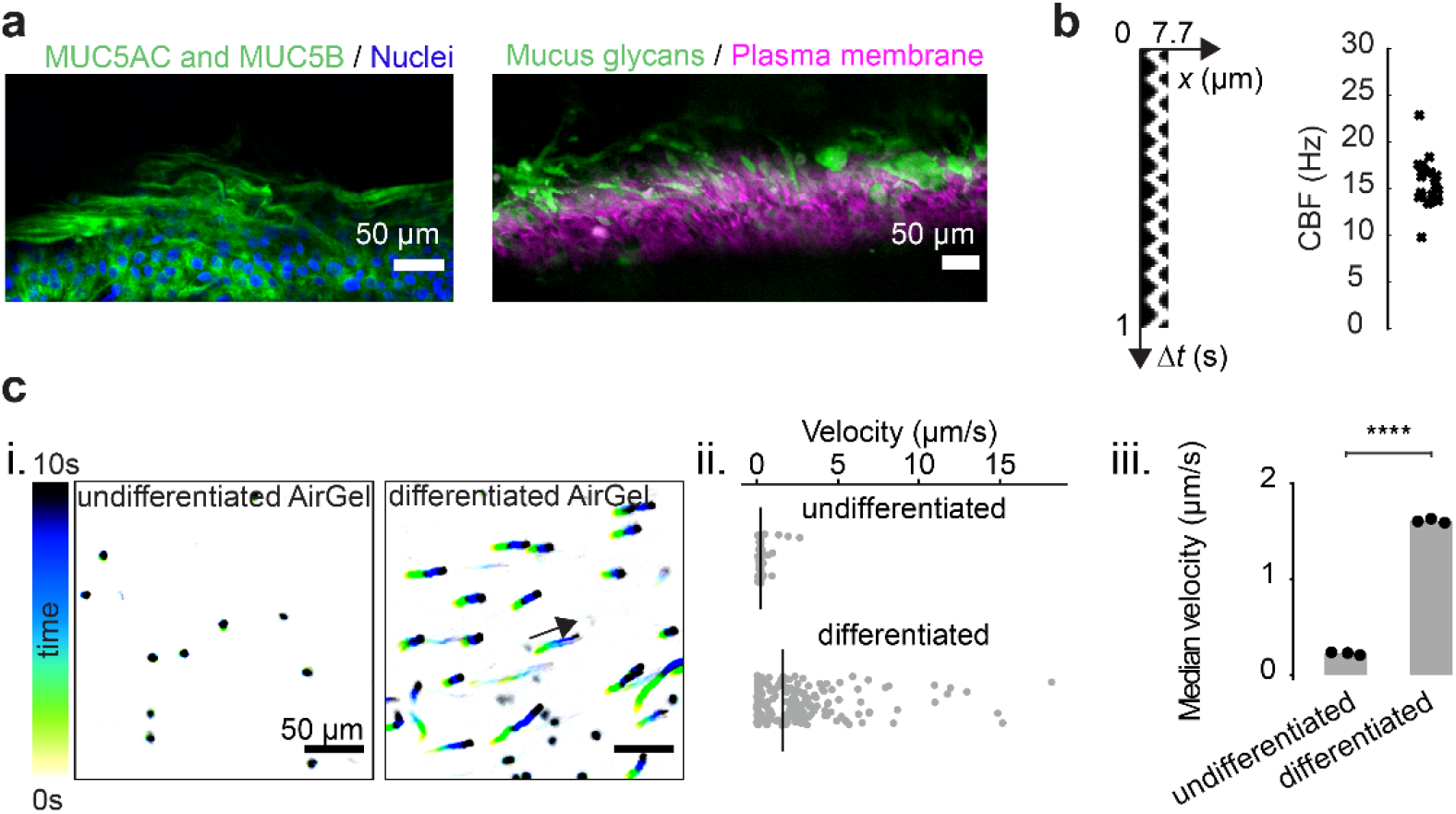
Mucociliary function of AirGels. **(a)** Extracellular luminal mucus in AirGels in methacarn-fixed (left) and live configurations (right). Stainings were done with antibodies against MUC5AC and MUC5B gel-forming mucins, as well as the fluorescent lectin jacalin (which targets glycans), respectively. **(b)** Cilia beating frequencies of 5 different AirGels, measured by tracking the oscillations of fluorescent beads attached to cilia. The kymograph on the left shows the trajectory of such a bead during one second. **(c)** Mucociliary clearance in AirGels. i. trajectories of fluorescent microparticles in the lumen of one undifferentiated and one differentiated AirGel. ii. corresponding velocity distributions. Black lines indicate the median velocity. iii. median particle velocities for three differentiated and undifferentiated AirGels show the contribution of cilia beating in clearance. Each data point corresponds to the median in each experiment; the gray bar shows the median of triplicates. Statistics: independent samples Student t-test with Bonferroni correction (*p* < 10^−7^).

### *P. aeruginosa* rapidly forms mucus-associated biofilms in AirGels

To visualize biofilm formation in a realistic airway mucosal context, we inoculated *P. aeruginosa* constitutively expressing the fluorescent protein mScarlet in the lumen of AirGels maintained at the air-liquid interface. After 13h of incubation, we observed that bacteria had extensively colonized the mucosal surface. *P. aeruginosa* formed interconnected bacterial clusters colocalized with mucus within the airway surface liquid (ASL) between epithelial cells and the air-liquid interface (Figure 3a). In dynamic visualizations, bacteria remained attached to mucus despite movements induced by beating cilia (Supplementary Video 2). Since *P. aeruginosa* takes days to form biofilms *in vitro*, we were surprised to see these communities form only within a few hours in AirGels^47^. We therefore went on to investigate the mechanisms of biofilm formation on mucus.

**Figure 3:**
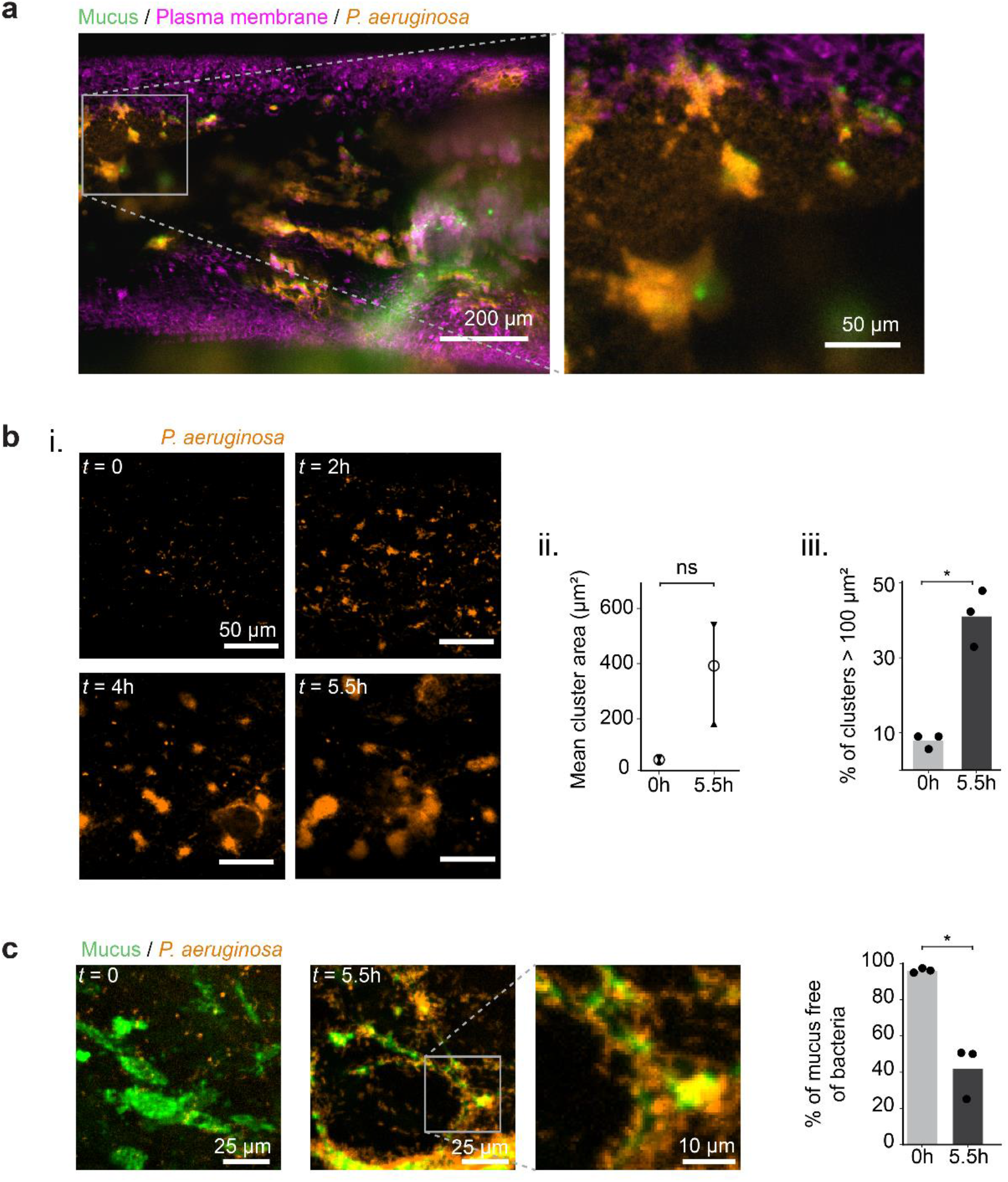
*P. aeruginosa* rapidly forms mucus-associated biofilms. **(a)** *P. aeruginosa* infection of a 62-day-old AirGel. Confocal images were acquired 13h after inoculation. *P. aeruginosa* constitutively expresses the fluorescent protein mScarlet. The plasma membrane of epithelial cells was stained with CellMask Deep Red (pink). Mucus was stained with jacalin (green) shortly before infection. **(b)** i. maximal intensity projection images show *P. aeruginosa* biofilm formation within hours. ii. mean biofilms cluster area for three AirGels. The bar indicates the range between the maximum and minimum of the three means. The circle represents the mean of the means. iii. percentage of clusters that were larger than 100 µm^2^ in each replicate (black dots). The bars represent the mean across replicates. Statistics: paired samples Student t-test with Bonferroni correction (*p* = 0.051 and *p* = 0.01). **(c)** *P. aeruginosa* rapidly colonizes mucus surfaces. Images show maximal intensity projection of confocal stacks at *t* = 0 and *t* = 5.5h post-inoculation. The graph quantifies the proportion of mucus not occupied by bacteria. Gray bars show the mean of triplicates. Statistics: paired samples Student t-test with Bonferroni correction (*p* = 0.02).

We imaged biofilm biogenesis in AirGels at the single cell level using confocal spinning disk microscopy. *P. aeruginosa* already formed aggregates a few hours after inoculation (Figure 3b). While the mucus surface was initially largely devoid of bacteria, half of it was covered by *P. aeruginosa* multicellular structures after 5.5h of infection (Figure 3c). Bacterial clusters with the same architecture also formed in the absence of jacalin staining, confirming these biofilms do not form through labeling artefacts (Supplementary Figure 2b). To confirm the pivotal role of mucus in biofilm formation, we infected a non-differentiated AirGel which does not produce mucus. In the absence of a protective mucus layer, epithelial cells were more vulnerable to *P. aeruginosa* infection (Supplementary Figure 3). Bacteria breached through the epithelial barrier and invaded the underlying ECM. *P. aeruginosa* did not form three-dimensional multicellular structures in the ASL. This further demonstrates the role of mucus hydrogel as a substrate for biofilm formation in differentiated AirGels, and at the same time highlights its protective function.

Our data suggests that *P. aeruginosa* forms biofilms in the airway by attaching to mucus at early stages of infection. To further explore the biophysical mechanisms of biofilm formation, we harvested mucus to perform *ex situ* visualizations. However, we could not observe the formation of *P. aeruginosa* biofilms on mucus extracted from HBE cultures (Supplementary Figure 4 & Supplementary Video 3). We attribute this discrepancy to perturbations in mucus mechanical integrity when extracted from the epithelium and immersed in buffer. This difference highlights the importance of investigating microbe-mucus interactions in a native mucosal context such as the one established in AirGels.

To understand how biofilms form on native mucus, we therefore inspected the different steps of their formation in AirGels. To nucleate *in vitro* biofilms, *P. aeruginosa* cells navigate the surface of abiotic materials using twitching motility, which promotes the formation of aggregates^46^. Fast imaging of single cells shows that *P. aeruginosa* moves with twitching-like trajectories at the surface of mucus fibers (Supplementary Video 4). Single cells aggregated into small clusters within 2h of colonization (Figure 3b). These small multicellular clusters subsequently moved along mucus fibers to eventually fuse into larger biofilms (Figure 4a). This caused a cascade of cluster fusion events that sped up biofilm formation (Figure 4a & Supplementary Video 5). We tracked aggregate fusion in kymographs highlighting the correlation between mucus and bacterial displacements (Figure 4b). The size of individual clusters remains approximately constant during motion and fusion, showing aggregate fusion predominates over bacterial growth. After 6h of aggregation and fusion, dense biofilms are formed.

**Figure 4:**
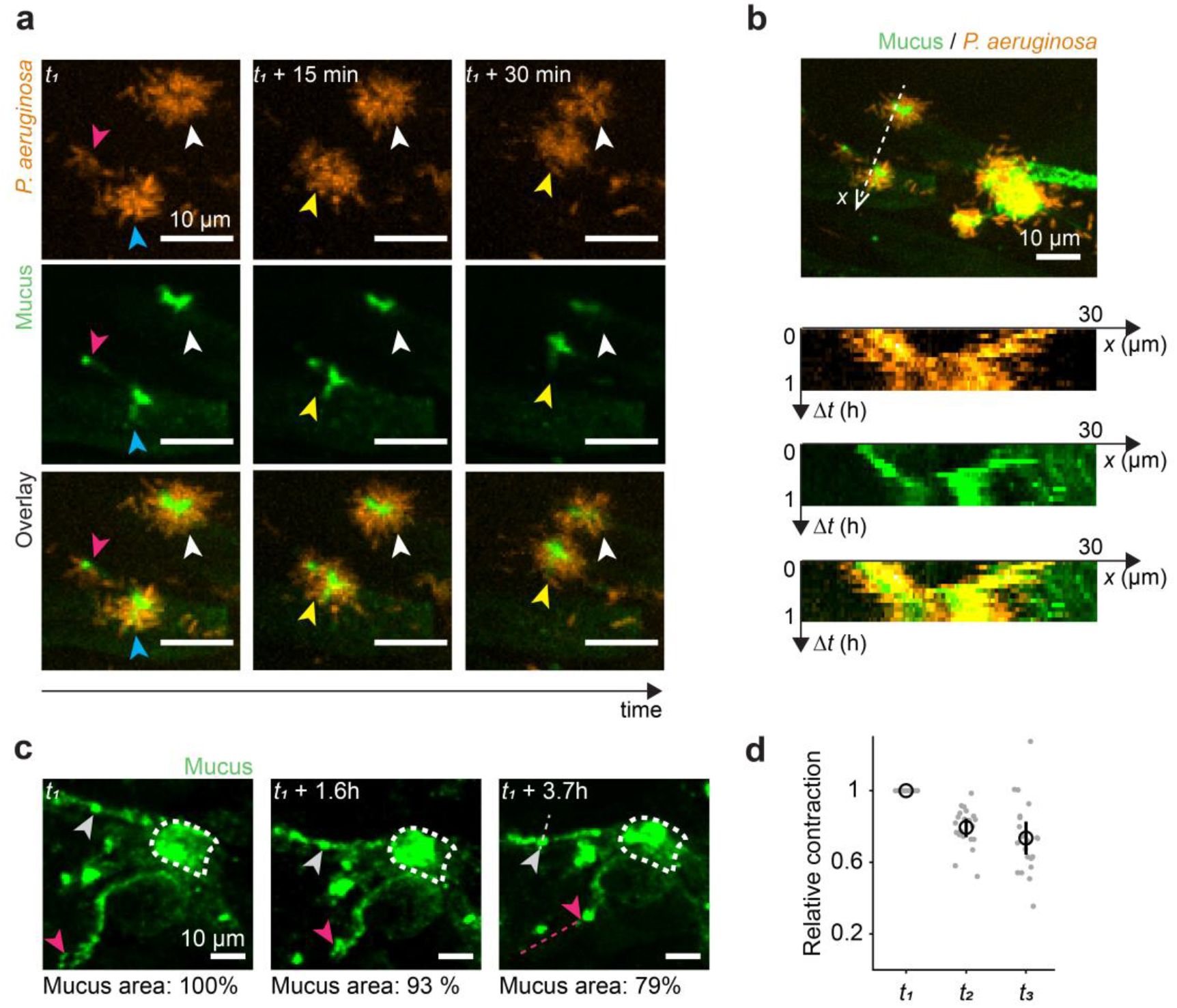
dynamics of biofilm formation on mucus. **(a)** Dynamic visualization of *P. aeruginosa* cluster fusion on mucus (*t*_*1*_ *=* 6.2h). The blue and pink arrowheads show two aggregates that fuse within the first 15 min. The resulting cluster is indicated by a yellow arrowhead. This new cluster then moves closer to the one indicated by the white arrowhead. All images are maximal intensity projections from z-stacks. **(b)** Kymographs showing the displacement of two clusters along their axis of motion. The bacterial aggregate and underlying mucus traveled together at an approximate speed of 0.5 µm/min. **(c)** Time course visualization of a mucus patch in a colonized by WT *P. aeruginosa* (not displayed). Reference features and their trajectories are indicated by colored arrowheads and dashed lines (*t*_*1*_ = 1.2h). **(d)** Mucus contraction was quantified by tracking the distances over time between *N =* 7 reference features in the mucus patch. The distances were normalized to the initial time point. They decrease over time, indicating contraction of the mucus patch. t_1_, t_2_, t_3_ refer to the timepoints shown in panel c. Black circle: mean. Black line: standard deviation.

We found that during biofilm formation, the mucus surface area tends to decrease compared to an uninfected control (Figure 4c & Supplementary Figure 5). The distances between landmarks in a mucus patch decreased over time (Figure 4d), demonstrating that mucus contracts during biofilm formation. We therefore hypothesized that mucus contraction speeds up biofilm formation by bringing *P. aeruginosa* cells closer to each other. Ultimately, these cells would become contiguous to form small aggregates. By carrying on mucus contraction, these aggregates would then fuse to each other. To substantiate this physical contraction mechanism, we investigated how *P. aeruginosa* could restructure mucus during attachment and colonization.

**Figure 5:**
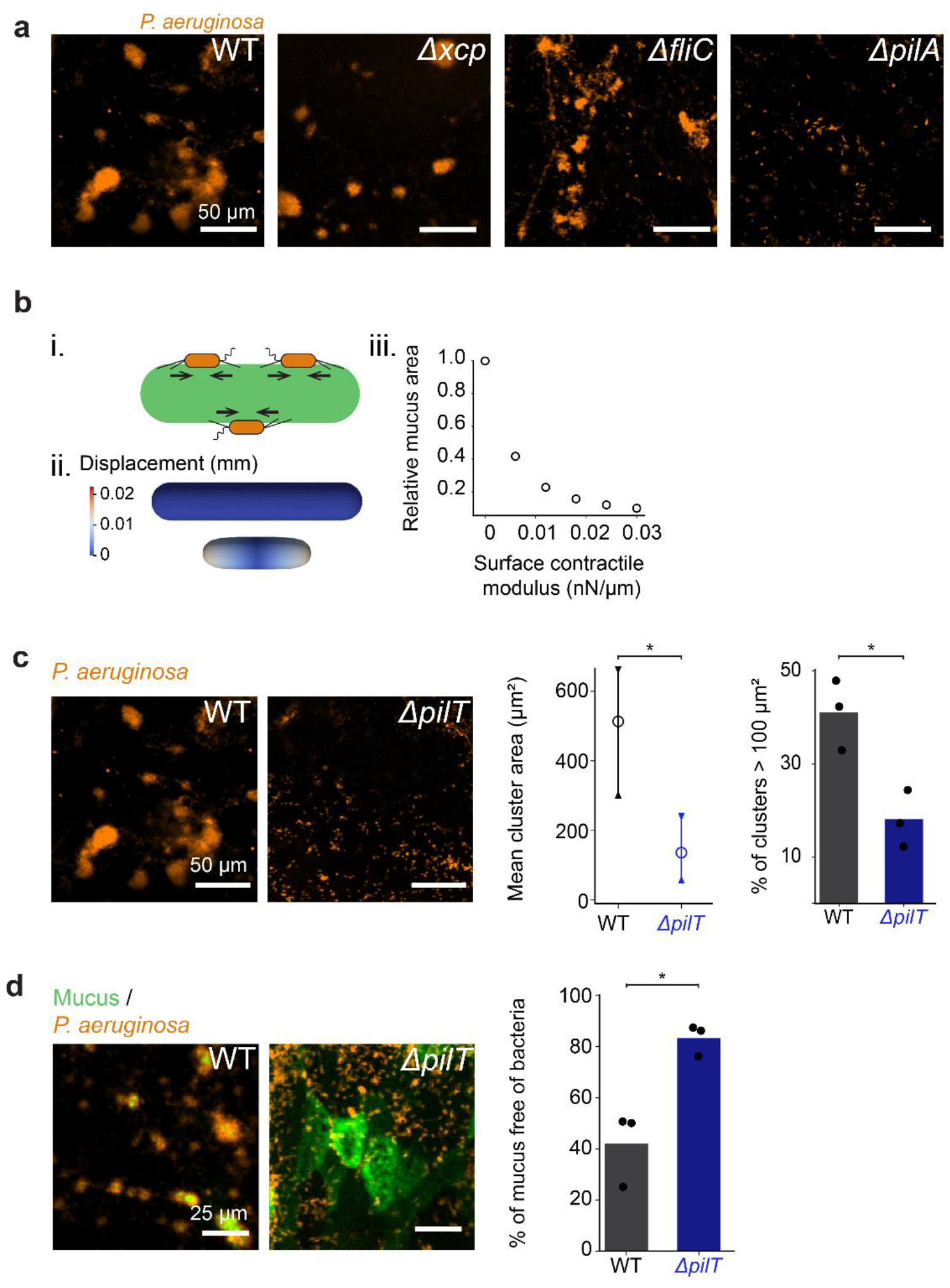
Type IV pili retraction promotes mucus contraction. **(a)** Biofilm formation of PAO1 mutants unable to degrade mucus or to generate force (*t* = 5.5h). Both the *Δxcp* mutant (lacking type II secretion system necessary for secretion of mucinases) and the *ΔfliC* mutant (lacking flagella) form biofilms similar to WT. In contrast, the *ΔpilA* mutant lacking T4P was unable to form biofilms in AirGels. **(b)** Finite element simulations of mucus deformation during surface contraction. i. schematic representation of *P. aeruginosa* applying contractile force on mucus by retracting T4P. ii. finite element simulation of deformation of a mucus cylinder at rest (top) and under active surface stress (bottom). Colormap indicate displacement of surface elements. iii. relative mucus area as a function of surface contractile modulus. As the surface contractile modulus increases, the relative area of mucus decreases. **(c)** T4P retraction is necessary for biofilm formation. Images compare biofilms from WT *P. aeruginosa* and from the *ΔpilT* mutant unable to retract T4P (*t* = 5.5h). *ΔpilT* cluster area and percentage of large clusters is significantly smaller than WT (*N* = 3). Statistics: independent samples Student t-test with Bonferroni correction (*p* = 0.035 and *p* = 0.015). **(d)** Mucus does not rearrange during *ΔpilT* colonization (*t* = 5.5h). Most of the mucus surface remains free of bacteria during *ΔpilT* colonization (*N* = 3). Statistics: independent samples Student t-test with Bonferroni correction (*p =* 0.01).

### *P. aeruginosa* forms biofilms on mucus using T4P

We envisioned two mechanisms for bacteria-induced mucus deformations: degradation or direct mechanical contraction. *P. aeruginosa* secretes mucinases capable of degrading gel-forming mucins^47^. Enzymatic mucus degradation could release polymers that generate entropic depletion forces promoting bacterial aggregation or that generate osmotic forces compressing mucus^14,48^. To test whether mucus degradation could drive contraction, we colonized AirGels with a mutant in the type II secretion system locus *xcp* that is necessary for mucin utilization^47,49^. The *Δxcp* mutant however formed biofilms similar to WT, ruling out the hypothesis of polymer-induced forces driving the formation of multicellular structures (Figure 5a).

Could *P. aeruginosa* remodel mucus by directly and actively applying force on the surface? *P. aeruginosa* can generate extracellular forces using flagella and T4P, motorized filaments that also play a role during *in vitro* biofilm biogenesis. In addition, T4P and flagella mediate single cell interactions with mucins^12,15,50–52^. To investigate their functions in the context of biofilm formation on mucus, we infected AirGels with *P. aeruginosa* mutants lacking flagella (*ΔfliC*) and T4P (*ΔpilA*). The *ΔfliC* mutant formed biofilms that were indistinguishable from wild type (WT) (Figure 5a). By contrast, *ΔpilA* cells did not form multicellular structures, indicating T4P play a role in mucus-associated biofilm formation. Since T4P may bind to glycans present on mucins^51,52^, weaker cell attachment to mucus could cause a decrease in aggregation of *ΔpilA*. Yet, colocalization shows that the *ΔpilA* mutant is still able to attach efficiently to mucus (Supplementary Figure 6 & Supplementary Video 6). We therefore envisioned a mechanism where T4P generate retractile forces that contract mucus from the surface, ultimately speeding up *P. aeruginosa* aggregation and cluster fusion.

To physically explore this scenario, we ran non-linear finite element simulations wherein mucus is treated as a hyperelastic material^53^. The mechanical action of *P. aeruginosa* T4P at the mucus surface is considered through the introduction of an active surface stress. The simulations recapitulated the experimental observations of mucus hydrogel contraction during *P. aeruginosa* colonization (Figure 5b). Simulations also predict that the steady-state mucus area decreases with the magnitude of the surface contractile modulus. This suggests that the more T4P retract, the more *P. aeruginosa* compresses mucus. To experimentally validate this model, we visualized AirGels colonization by a *ΔpilT* mutant which produces T4P that cannot retract, mimicking conditions of zero contractile modulus. *P. aeruginosa ΔpilT* could still associate with mucus and form a few small clusters, but clearly failed to form mucus-associated biofilms to the same extent as WT (Figures 5c & d), which was coherent with simulations. These results show that T4P retraction is necessary for biofilm formation on mucus, and is consistent with a scenario where retraction compresses the mucus substrate.

To further support the surface contraction model, we tested the prediction that deformations increase further with surface contractility. We imaged AirGel colonization by the hyperpiliated *P. aeruginosa* mutant *ΔpilH*, whose T4P retraction frequency is approximately twice the one of WT (Supplementary Figure 7). *ΔpilH* formed biofilms more rapidly than WT: we observed dense aggregates as early as 2h, while we only did after 4h for WT (Figure 6a & b). In addition, *ΔpilH* induced more rapid mucus contraction than WT (Figure 6c & Supplementary Videos 7 & 8), consistent with simulations. After 5.5h, WT and *ΔpilH* biofilms had similar morphologies and size, suggesting biofilm fusion reaches a physical limit most likely due to packing at the mucus surface. To control that the observed differences did not arise from growth rate variation between the strains, we quantified fluorescence intensity over time for WT, *ΔpilT* and *ΔpilH*. All strains appeared to grow at similar rates in the ASL of AirGels (Supplementary Figure 8).

**Figure 6:**
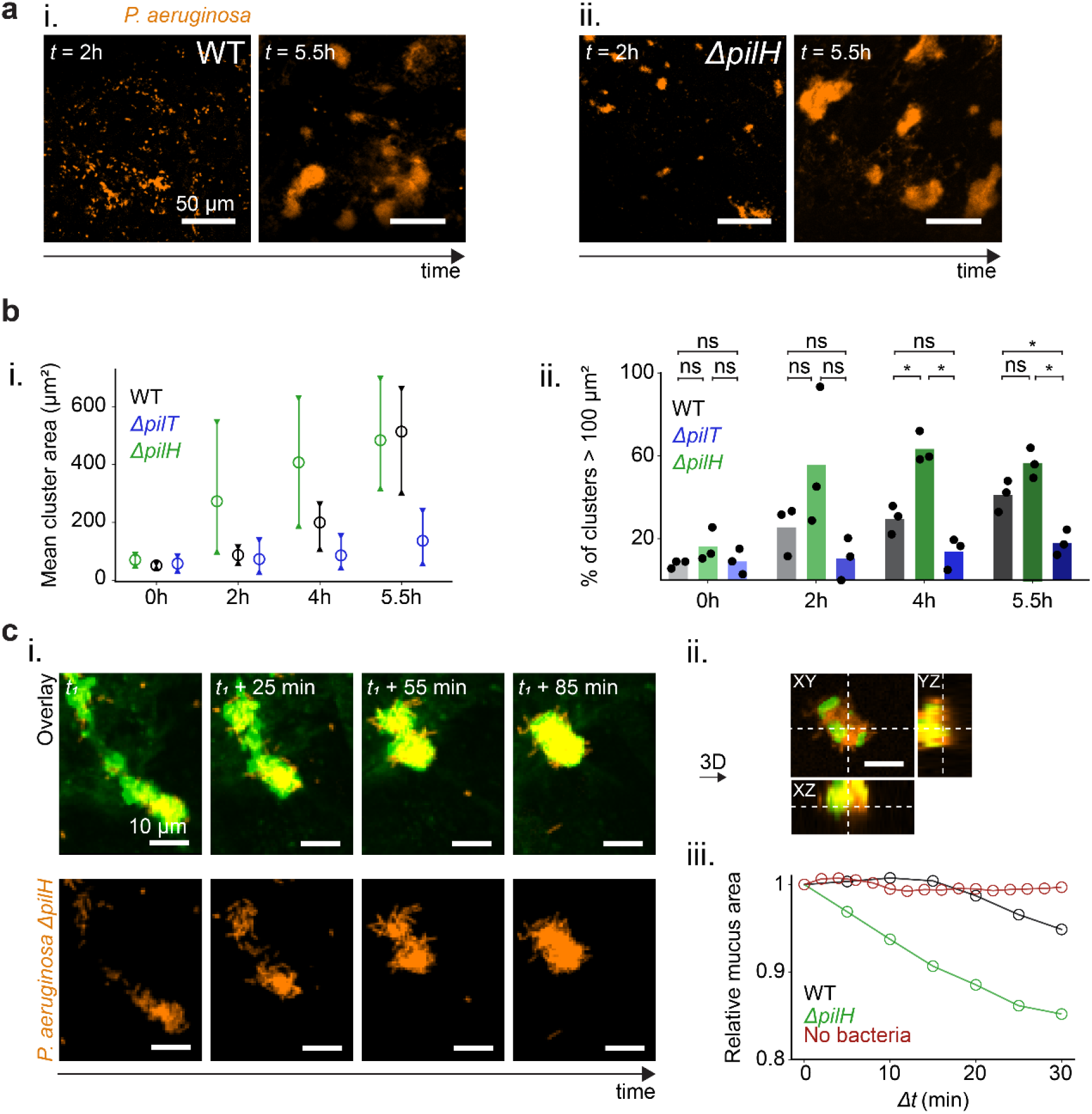
Increased T4P retraction speeds up biofilm formation and mucus contraction. **(a)** Increased T4P activity speeds up biofilm formation on mucus. Comparison of biofilm formation by the *ΔpilH* mutant with hyperactive T4P versus WT, at *t* = 2h and *t* = 5.5h after inoculation. *ΔpilH* already forms small biofilms after 2h. **(b)** i. kinetics of biofilm size for WT, *ΔpilT* and *ΔpilH*. For each strain, we infected three AirGels from a healthy donor. Bars represent the range between the maximum and minimum of the means from triplicates, circles represent the overall mean. ii. comparison of percentage of large clusters for WT, *ΔpilT* and *ΔpilH*, over time. Statistics: one-way ANOVA for each time point, followed by a post-hoc Tukey test if the null hypothesis was rejected. At *t* = 4h, the differences between WT and *ΔpilH* (*p* = 0.003) and between *ΔpilH* and *ΔpilT* (*p* = 0.001) were significant. At *t* = 5.5h, the differences between WT and *ΔpilT* (*p* = 0.02) and between *ΔpilH* and *ΔpilT* (*p* = 0.001) were significant. **(c)** *ΔpilH* dramatically contracts mucus. i. timelapse images showing an event of mucus contraction by *ΔpilH*. ii. orthogonal views of the bacteria-covered mucus cluster at *t*_*1*_ + 85 min, showing that PAO1 *ΔpilH* cells surround mucus. iii. relative mucus area changes measured during 30 min for WT and *ΔpilH*; since *ΔpilH* starts aggregating and remodeling mucus earlier than WT, the starting points of the recording differed (*ΔpilH*: 2.5h, WT: 6.2h, negative control: 8.1h). Images are maximal intensity projections of z-stacks throughout the figure except for the orthogonal projection in H.

Overall, our results support a model where *P. aeruginosa* contracts the surface of mucus by active T4P retraction. Single cells initially twitch on mucus to form small aggregates. The static aggregate collectives generate forces from T4P that are sufficient to deform their substrate, driving large-scale mucus contraction. By contracting, mucus brings aggregates closer. They eventually fuse and form biofilms (Figure 7).

**Figure 7:**
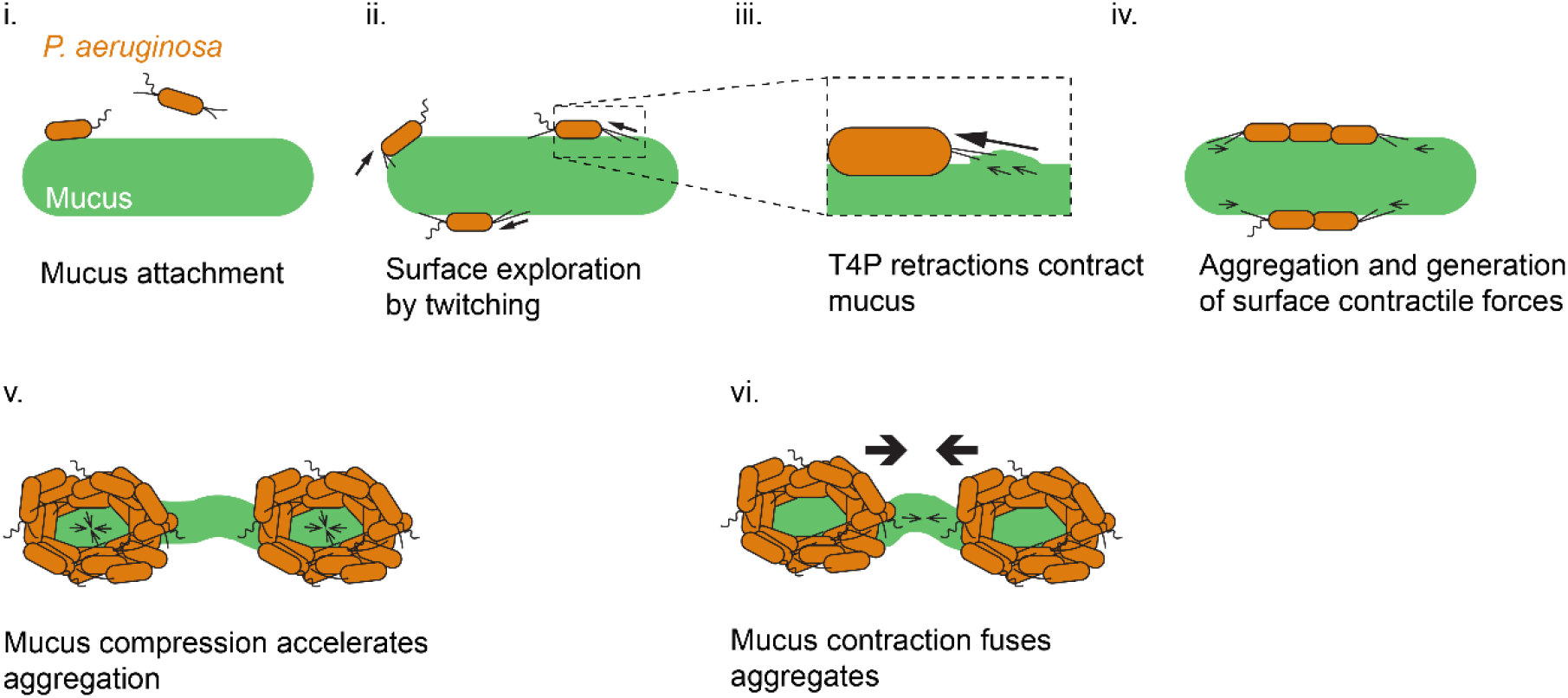
Proposed model for the formation of mucus-associated biofilms by *P. aeruginosa*. i) Single bacterial cells attach to mucus. ii) *P. aeruginosa* navigates the mucus surface using T4P-dependent twitching motility. iii) T4P retraction locally contracts mucus. iv) Surface exploration promotes encounters between single cells. This initiates aggregation and the formation of small clusters. These clusters remain static on mucus. As single cells in aggregates use T4P to pull on and contract mucus, they generate surface contractile forces. v) Collective retraction of T4P from many cells compacts bacterial aggregates. vi) Still under the action of retractile T4P forces, aggregates further contract mucus to initiate fusion into biofilms.

## DISCUSSION

We used AirGels, tissue-engineered human lung organoids, to investigate how *P. aeruginosa* forms biofilms at the airway mucosal surface. We found that *P. aeruginosa* forms biofilms via an active mechanism of mucus remodeling. *P. aeruginosa* attaches to mucus and subsequently uses T4P to generate surface contractile forces. As a result, the mucus gel contracts, effectively reducing its area and bringing mucus-bound bacteria closer to each other. After this initial biofilm seeding, *P. aeruginosa* may initiate the secretion of matrix components to strengthen the cohesion of the biofilm.

T4P play multiple functions during biofilm formation in many species. In *P. aeruginosa*, successive T4P extension and retraction power twitching motility on surfaces^54,55^. This allows freshly attached cells to explore the environment, stimulating cell-cell encounters that nucleate aggregation^55^. These microcolonies ultimately mature into full biofilms. This model however falls short on soft surfaces. Hydrogels with low stiffness limit the transmission of T4P traction force to the surface thereby impairing twitching motility, but at the same time still enable biofilm formation^56^. Mucus contractions induced by *P. aeruginosa* show that T4P-generated forces can remodel soft materials as well. In addition to highlighting a new mode of biofilm formation, this mechanism provides additional evidence that bacteria can mechanically remodel the host microenvironment^57^.

Although the classical view of airway infections associates biofilms with chronic infections and planktonic cells with acute infections, recent work has demonstrated the coexistence of these bacterial lifestyles in sputum samples from both acutely and chronically infected patients^7^. Our observations of early biofilm formation in AirGels is therefore consistent with these clinical observations. However, *in vitro*, the effect of mucus on biofilm formation depends on the experimental protocol: while native mucins in solution inhibit biofilm formation^15–17^, studies with full mucus or commercial mucins instead report increased aggregation^11,12,14^. This demonstrates that mucus-pathogen interactions vary dramatically depending on the model system used, thereby highlighting the importance of carefully reproducing and controlling relevant parameters *in vitro*.

Forming biofilms early on could provide a fitness advantage to *P. aeruginosa* in the non-hospitable airway environment. For example, bacterial aggregation could reduce *P. aeruginosa*’s susceptibility to neutrophils which are rapidly recruited to the mucosal surface during infection^58,59^. At the same time, forming biofilms increases *P. aeruginosa* tolerance to antibiotic treatment and promotes the emergence of resistant mutants^60^. There is however an upside for the host: mucus adsorbs a large proportion of the planktonic *P. aeruginosa*, keeping them away from the epithelium. Our results therefore highlight the duality of mucus: protecting the airway epithelium from acute infections, while providing a fertile ground for biofilm formation that favors chronic infections.

Most investigations of host-pathogen interactions have so far mainly relied on animal models and immortalized cell lines. Their limitations have been an obstacle to establish a holistic understanding of infections. By developing AirGels, we provide the community with a 3D airway infection model that expresses relevant cell types, secretes mucus, and is compatible with high-resolution imaging in presence of an air-liquid interface. As a result, AirGels have a strong potential in bridging the gap between *in vivo* and *in vitro* investigations of airway infections. Since AirGels are modular, we envision that engineering refinements could improve their suitability as an infection model for a wide range of organisms. In addition, characterizing the rheological and chemical properties of the mucus lining would allow a better connection between investigations in AirGels and clinical observations. In summary, alternative approaches that leverage engineered microenvironments will help us better comprehend bacterial physiology in realistic infection contexts, which could ultimately allow the discovery of novel therapeutic strategies against antibiotic-resistant infections.

## ACKNOWLEDGEMENTS

We thank Zaïnebe Al-Mayyah, Jeremy Wong, the Bioimaging and Optics Core Facility (BIOP) and the Gene Expression Core Facility (GECF) at EPFL for technical assistance. Marco Kühn, Johannes Bues and Sophia Hsin-Jung Li for insightful discussions, Romé Voulhoux for strains and Formlabs forum user Telliria for suggestions on 3D-printing process. We acknowledge the Swiss National Science Foundation for funding this project through the Project grant number 310030_189084 and NCCR AntiResist.

