## Supplementary material for "*Pseudomonas aeruginosa* contracts mucus to form biofilms in tissue-engineered human airways"

### Supplementary material for:
### *Pseudomonas aeruginosa* contracts mucus to rapidly form biofilms in tissue-engineered human airways

**Authors:**

Tamara Rossy[1,2], Tania Distler[1,2], Joern Pezoldt[2], Jaemin Kim[3], Lorenzo Talà[1,2], Nikolaos Bouklas[3], Bart Deplancke[2], Alexandre Persat[1,2]

**Affiliations:**

[1] Global Health Institute, School of Life Sciences, Ecole Polytechnique Fédérale de Lausanne (EPFL), Lausanne, Switzerland

[2] Institute of Bioengineering, School of Life Sciences, École Polytechnique Fédérale de Lausanne (EPFL), Lausanne, Switzerland

[3] Sibley School of Mechanical and Aerospace Engineering, Cornell University, Ithaca, NY, USA

### Supplementary figures & captions

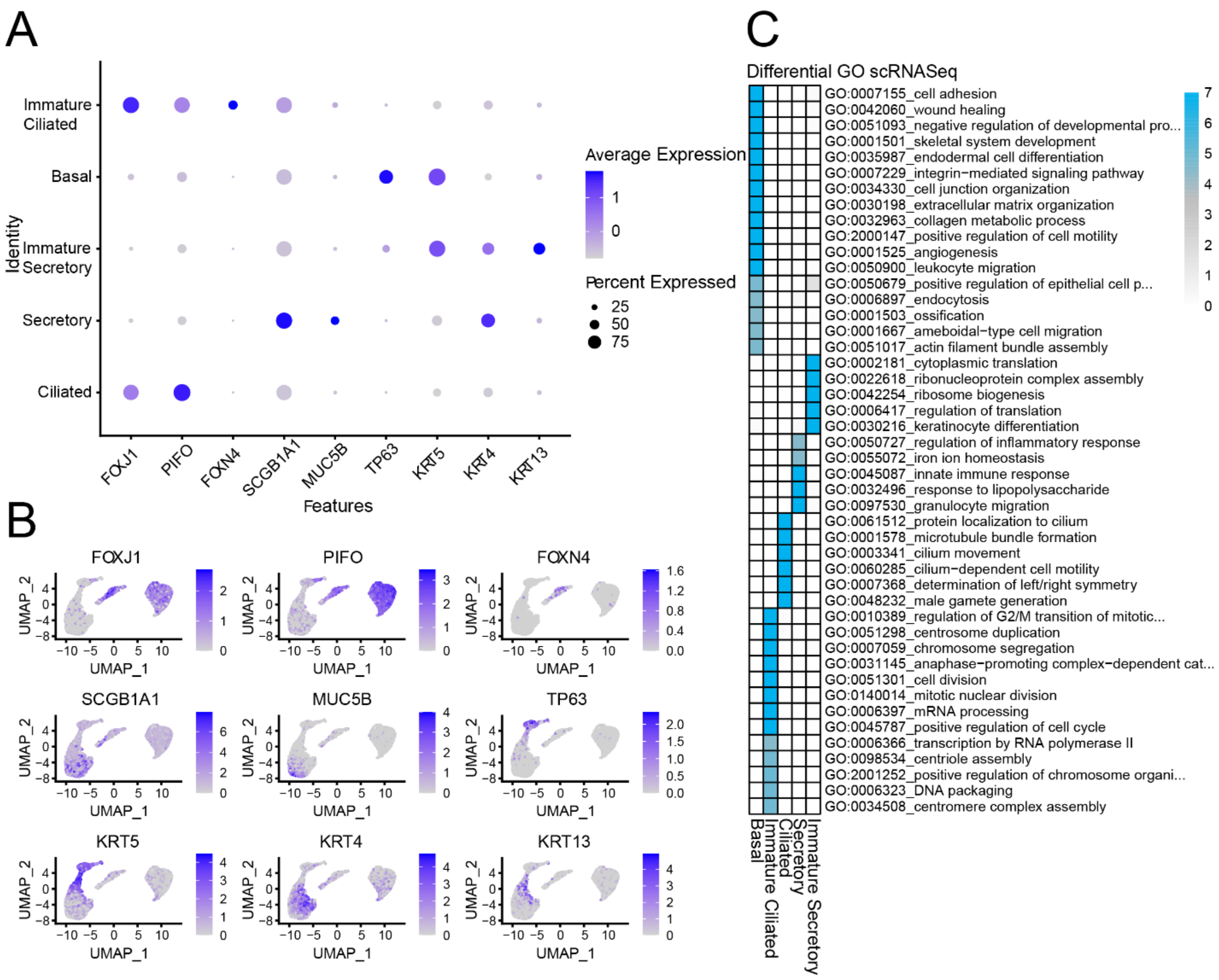


**Supplementary Figure 1:** Gene expression profiles of AirGels measured by single cell RNA-seq. **A.** The expression of marker genes of lung epithelial cell types is shown for each cluster defined from the scRNA-seq reads of AirGels. Average expression per cluster and the percentage of cells expressing the respective marker gene. **B.** Subset-specific expression of canonical marker genes on UMAP embedding. FOXJ1 and PIFO are typically expressed in ciliated cells. The immature ciliated cell cluster, also known as deuterosomal cells, is marked by high levels of FOXJ1 and expression of FOXN4. Basal cells typically express TP63 and KRT5. The secretory cluster shows expression of SCGB1A1 and a fraction of more mature secretory cells expressing MUC5B[1]. Furthermore, we observe a transitional state between basal and secretory, the immature secretory cluster, which shows partial mutual expression of KRT4 and KRT13 as previously described[2]. **C.** A gene ontology (GO) analysis was performed on the most differentially-expressed genes in each cluster confirming the correct annotation of cell clusters.

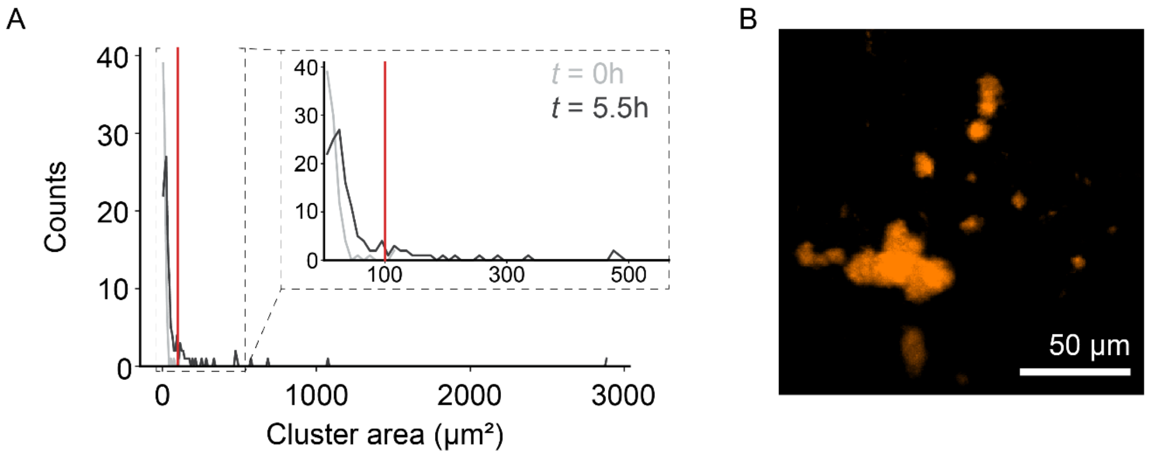


**Supplementary Figure 2:** Distributions of cluster sizes and unlabeled control for AirGels infected with *P. aeruginosa*. **A.** Distribution of cluster areas at $t$ = 0h and $t$ = 5.5h during one infection experiment. The red line indicates the threshold for what we considered as large clusters (> 100 µm$^2$). **B.** Infection of an AirGel in which mucus was not labeled. After 4h15, bacterial aggregates were already visible, indicating their formation is independent of jacalin staining.

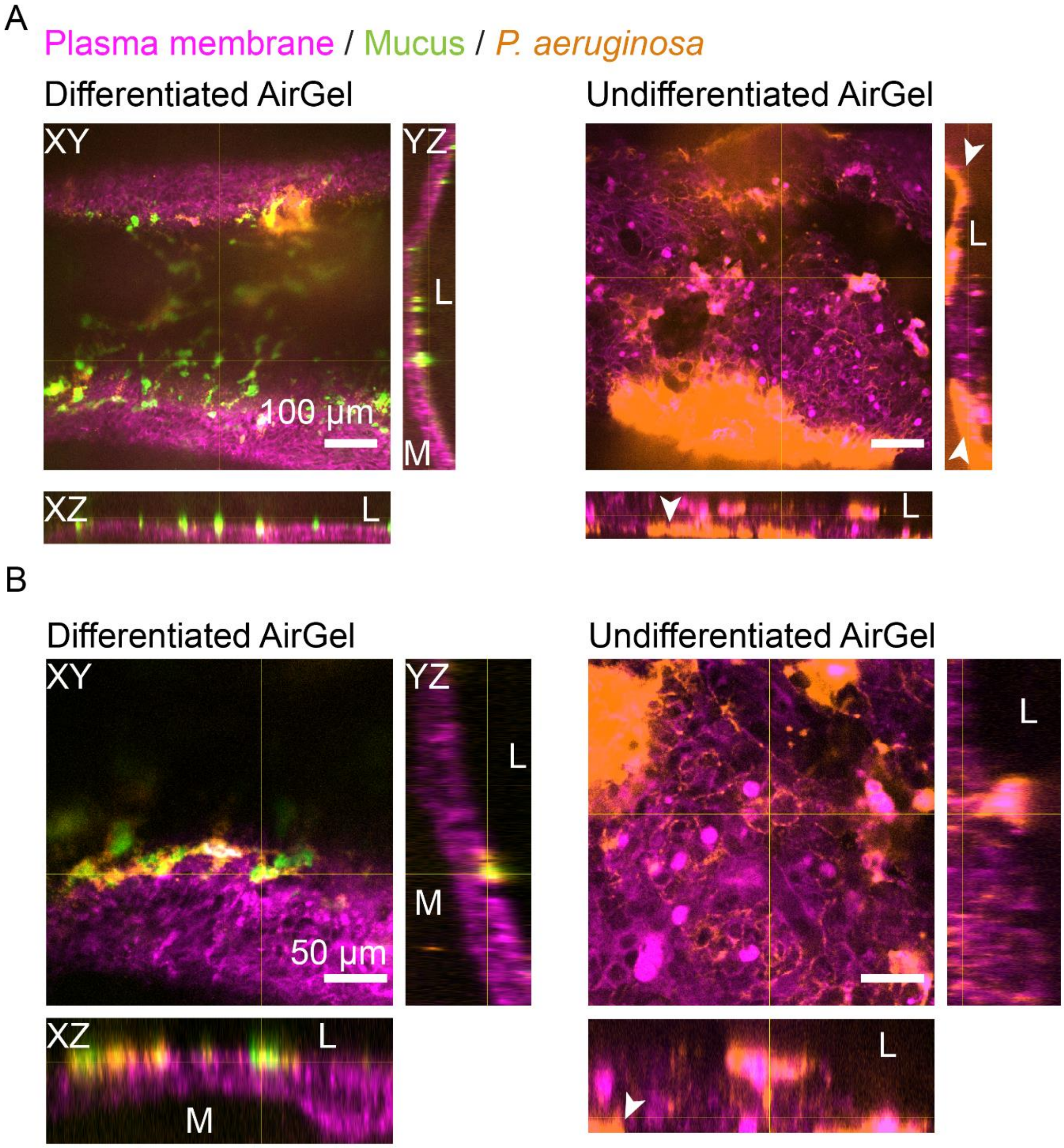


**Supplementary Figure 3:** *P. aeruginosa* does not form biofilms in AirGels lacking mucus, but damages its tissue more rapidly. Orthogonal views of infections with *P. aeruginosa* in a differentiated versus an undifferentiated AirGel stained with Jacalin at low **(A)** and high **(B)** magnification. The infection was imaged at $t$ = 5.5h (differentiated) and $t$ = 6h (undifferentiated) post-inoculation. L indicates the luminal side and M the extracellular matrix. We did not observe *P. aeruginosa* aggregates on the luminal side of the undifferentiated AirGel. However, bacteria damaged the epithelium extensively in the absence of mucus, which resulted in invasion of the extracellular matrix (white arrowheads).

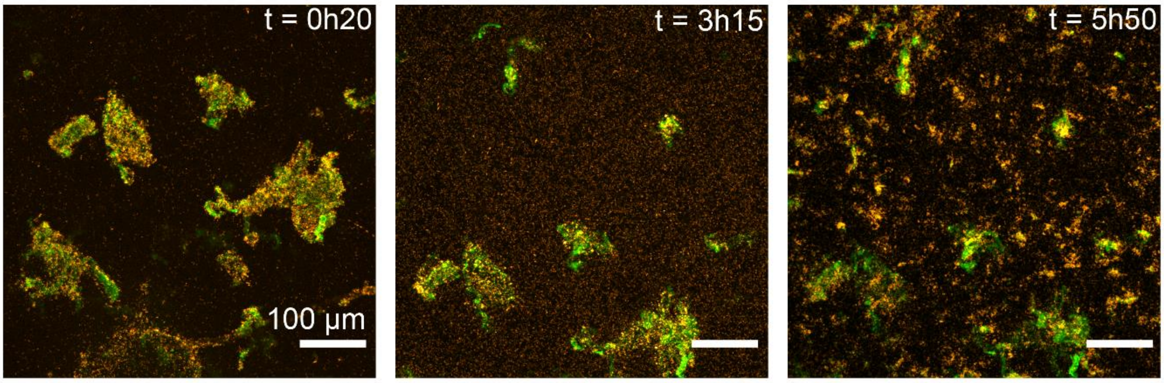


**Supplementary Figure 4:** *P. aeruginosa* does not form biofilms on mucus extracted from HBE cultures. Jacalin-labeled mucus that had been isolated from a differentiated HBE culture on a Transwell. Even after almost 6h after incubation with *P. aeruginosa*, no aggregates were visible.

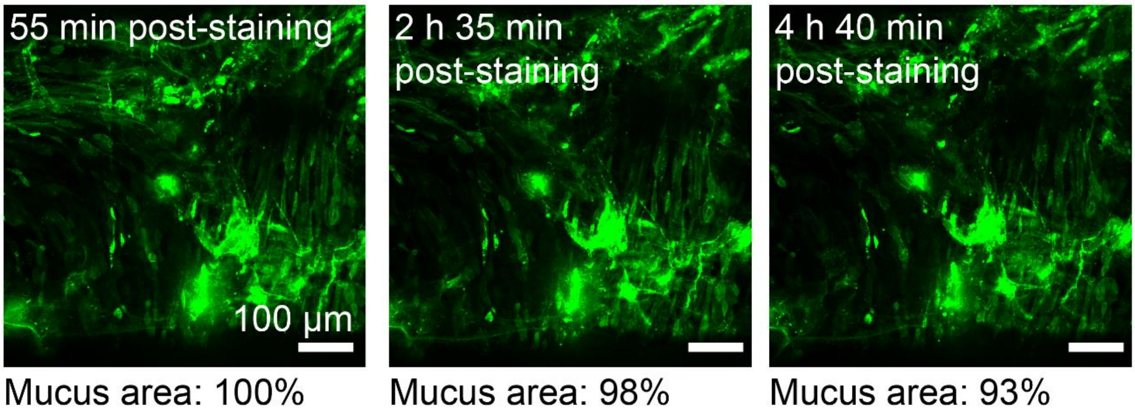


**Supplementary Figure 5:** Mucus does not contract in the absence of *P. aeruginosa*. Jacalin-stained mucus in an uninfected AirGel. The total area of mucus was estimated over time and found to only differ slightly over time, most likely due to photobleaching and drift in and out of focus.

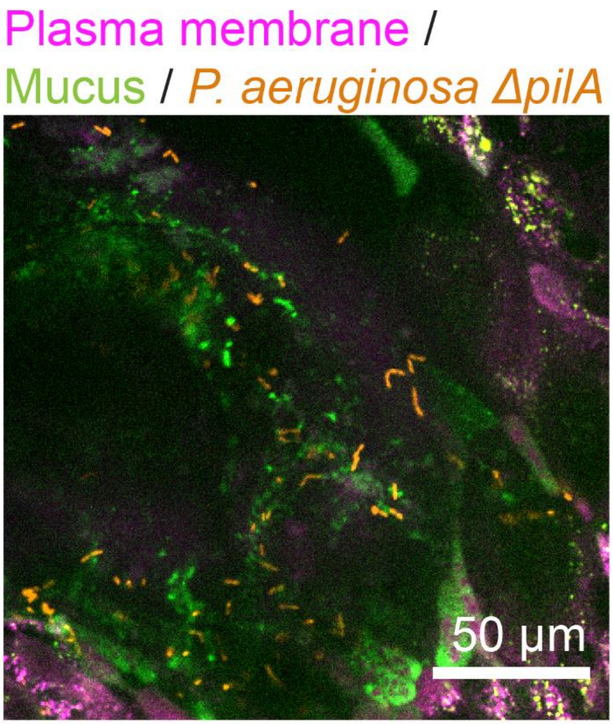


**Supplementary Figure 6:** T4P are necessary for mucus contraction but not for attachment. Aggregates of the *ΔpilA* mutant *P. aeruginosa* were still absent 3h25 post-inoculation. However, the bacteria colocalized with the mucus, indicating that T4P are not necessary for adhesion to mucus.

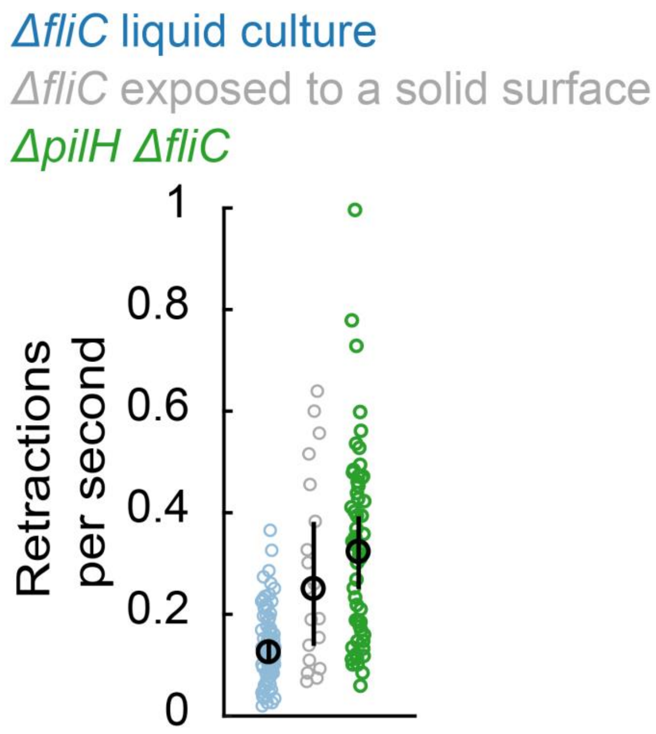


**Supplementary Figure 7:** T4P retraction frequency increases during surface contact and is constitutively high in a *pilH* deletion mutant. T4P retraction rates were measured by interferometric scattering (iSCAT) microscopy, which allows for label-free T4P visualization[3]. To prevent cells from swimming away during the iSCAT measurements, a flagellum-less *ΔfliC* mutant was used as background strain. This strain was either grown in liquid or pre-adapted to culture on a solid surface for 3h. Black circles and bars indicate the bootstrap median and 95% confidence interval of the medians, respectively.

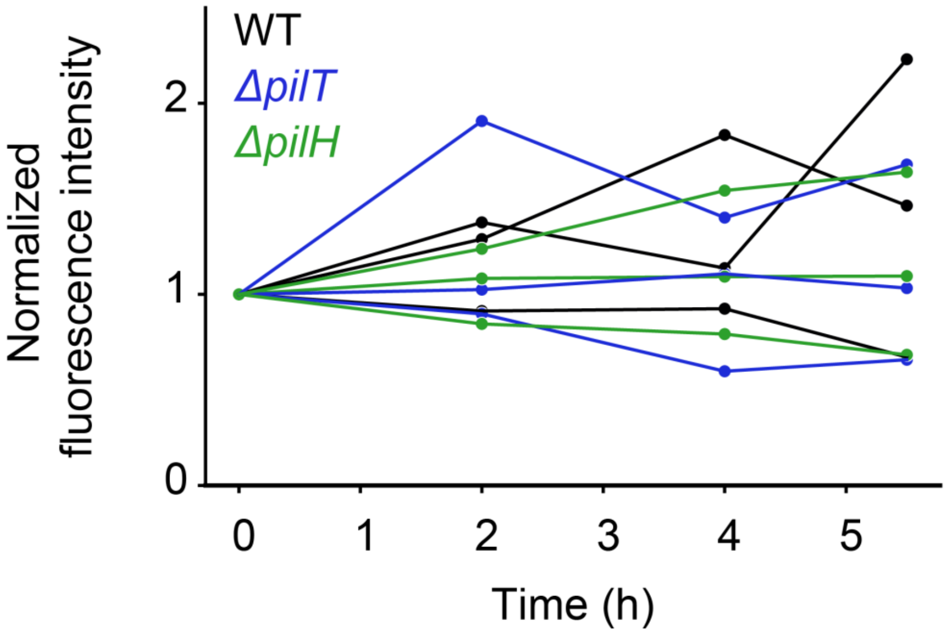


**Supplementary Figure 8:** Estimation of bacterial growth rate in $N$ = 9 AirGels for the different strains used in this study. Total fluorescence intensity summed across z-stack acquisitions was normalized to the $t$ = 0h data. All strains have indistinguishable growth rates at the ASL of AirGels.

### Captions for supplementary videos

**Supplementary Video 1:** real-time video recording of beating cilia in a differentiated AirGel.

**Supplementary Video 2:** network of mucus-associated *P. aeruginosa* biofilms 13h post-inoculation into an AirGel. Beating of the underlying ciliated epithelium shakes the biofilms, but is not sufficient to clear them out of the model airway.

**Supplementary Video 3:** time-lapse spinning disk confocal micrographs showing *P. aeruginosa* (orange) on jacalin-labeled mucus (green) isolated from a Transwell. Even 5h post-infection, the bacteria did not aggregate on nor remodel the mucus.

**Supplementary Video 4:** time-lapse spinning disk confocal micrographs of *P. aeruginosa* (orange) twitching on the mucus (green) of an HBE culture in a Transwell insert. The bacteria were loaded 3.5h before the time-lapse was recorded.

**Supplementary Video 5:** time-lapse spinning disk confocal micrographs of *P. aeruginosa* (orange) on jacalin-labeled mucus (green). The initial time stamp in the video corresponds to *t* = 6h10 post-inoculation. Several clusters can be seen fusing over the course of the experiment; large pieces of mucus also shrink upon interactions with bacteria.

**Supplementary Video 6:** time-lapse spinning disk confocal micrographs of a T4P-less *P. aeruginosa ΔpilA* (orange). The bacteria are moving along with the flowing, jacalin-labeled mucus (green), indicating that T4P are not necessary for adhesion to mucus.

**Supplementary Video 7:** compilation of 30 min time-lapses showing jacalin-labeled mucus (green) in different conditions: first, without any bacteria, then exposed to a WT *P. aeruginosa* strain (orange) and finally exposed to a hyperpiliated *ΔpilH* mutant. Since *ΔpilH* starts aggregating and remodeling mucus earlier than WT, the starting points of the recording differed (*ΔpilH*: 2 h 30 min, WT: 6 h 10 min, negative control: 8 h 5 min).

**Supplementary Video 8:** time-lapse spinning disk confocal micrographs of the hyperpilated *P. aeruginosa ΔpilH* mutant (orange) rearranging jacalin-labeled mucus (green). Dramatic remodeling of the mucus starts as early as 1.5h post-inoculation, resulting in numerous mucus-associated aggregates.

### Materials and methods

#### AirGel chip fabrication

##### 3D printed mold

The mold for the PDMS chips was designed in Autodesk Inventor Professional 2021. This mold was then 3D printed by Multi Jet Modeling on a Connex 500 printer (Objet) using VeroClear resin at the Additive Manufacturing Workshop (AFA) at EPFL. In order to remove uncured resin that could interfere with subsequent PDMS polymerization, we treated our mold by soaking it in deionized water for 2h; then, we incubated it for 18h in an oven set to 85°C; finally, we washed it with deionized water and dishwashing soap before letting it dry.

##### PDMS chip

PDMS (Sylgard 184, Dow Corning) was casted on the mold and cured at 60 °C for about 1h30. We then used a scalpel to cut out each chip individually and carefully remove it from the mold. In parallel, we prepared PDMS rods to pattern the lumens according to a published protocol [4]. In short, we filled gauge 14 needles (Sterican 2.1 x 80 mm, B. Braun) with PDMS and cured it as described above. We then used pliers to break the needles and extract the PDMS rods; their diameter was approximately 1.6 mm (i.e. inner diameter of the needle). We used a scalpel to cut them into 8mm-long pieces. Then, PDMS chips and rods were briefly immersed in isopropanol, left to dry, and cleaned using tape. Afterwards, the rods were inserted into the chips using tweezers; the assembled devices were subsequently autoclaved. They were then plasma bonded to either glass-bottom dishes or glass-bottom 6-well plates (1.5 coverslip, glass diameter 20 mm, MatTek) in a ZEPTO plasma cleaner (Diener electronic). Note that the chips contained thin PDMS membranes at the bottom of their inlet reservoirs (obtained owing to a shallow cavity in the 3D printed mold), so that the rod was not in direct contact with the underlying coverslip. Finally, we exposed the chips to two cycles of UV sterilization in a biosafety cabinet.

##### Extracellular matrix

All the following steps were performed in a biosafety cabinet to maintain sterility. We treated the chips inner surfaces to promote adhesion of the gel following a published method[5]. This consisted in a 10 min exposure to 2% polyethyleneimine (Sigma-Aldrich) followed by a 30 min treatment with 0.4% glutaraldehyde (Electron Microscopy Sciences) for 30 min. The chips were then rinsed once with Milli-Q water (Merck Millipore). Afterwards, the ECM hydrogel was prepared on ice. We first neutralized high-density rat tail type I collagen (~10 mg/ml, Corning) to a final concentration of 8 mg/ml. To do this, we mixed 30 µl 10X PBS, 5.4 µl NaOH 1M, 29.6 µl Milli-Q water and 235 µl collagen with a positive displacement pipette (Gilson). The neutralized collagen was then mixed with high-concentration growth factor reduced Matrigel (~21 mg/ml, Corning) in a 75:25 ratio (100 µl Matrigel for 300 µl of neutralized collagen). The resulting gel was loaded into each chip from the basal access ports and placed in a humidified cell culture incubator (set to 37°C, 5% $CO_2$) during 20 min in order for polymerization to occur. Then, we pulled the rods out of the chips with tweezers, thereby shaping the lumen[4]. The final step consisted in chemically crosslinking the collagen to strengthen it. We followed a published protocol[6]: we first prepared a 0.6M solution of N-(3-dimethylaminopropyl)-N′-ethylcarbodiimide hydrochloride (EDC, Life Technologies) and a 0.15M solution of N-hydroxysuccinimide (NHS, Sigma-Aldrich). Then, we mixed those solutions in a 1:1 ratio and loaded 25 µl into each lumen. We left them at room temperature for 5 min before aspirating the crosslinking reagents. We then soaked the chips in Milli-Q water (apical and basal sides) overnight, at room temperature. Finally, we replaced Milli-Q water with PneumaCult-Ex Plus medium (Stemcell Technologies) at least one day before loading any cells in the chips, and stored the chips in a cell culture incubator.

#### Cell culture

##### Expansion in flasks

We obtained primary human bronchial epithelial cells from Lonza (CC-2540S or CC-2540 for healthy donors, 00196979 for the CF donor). They were cultured in T-25 flasks using PneumaCult™-Ex Plus medium (Stemcell Technologies) for no more than 3 passages. When they reached confluence, the cells were detached from the flask using the Animal Component-Free Cell Dissociation Kit (Stemcell Technologies) and centrifuged before being resuspended in PneumaCult™-Ex Plus to a density of approximately 20’000 cells/µl.

##### Loading into AirGels

AirGels were emptied of all culture medium on the apical and basal sides. 10-12 µl of HBE cell suspension was loaded in the lumen of each AirGel. The chips were then placed in a cell culture incubator for 25 min, flipped upside down and incubated again for 25 min, and then finally for 15 min on each side in order to allow uniform adhesion of cells along the luminal surface. Afterwards, PneumaCult™-Ex Plus was added to the lumen and on the basal side of the chips.

##### Long-term culture in AirGels

HBE cells were expanded in AirGels with PneumaCult™-Ex Plus until confluence was reached (typically 1-3 days). After that, apical and basal expansion medium was replaced with PneumaCult™ Airway Organoid Differentiation Medium (Stemcell Technologies). In order to prevent gel degradation by the HBE cells, we supplemented the medium with 5 µM of protease inhibitor GM6001 (InSolution GM6001, Merck). One day after the first addition of differentiation medium, all fluid was aspirated from the lumen, thereby generating an air-liquid interface (ALI). This critical step was facilitated by the aforementioned collagen crosslinking and by the large lumen diameter, which lowered capillary forces drawing medium back in the channel. AirGels could be kept in these conditions for at least one month; medium was replaced on the basal side every second day. Before weekends, the lumen was also filled with PneumaCult™ Airway Organoid Differentiation Medium, but ALI was restored every Monday morning and maintained for the whole week.

##### Cell culture on Transwell membranes

For isolated mucus experiment and twitching motility imaging, we grew NHBE cells on 0.4µm-pore polyester Transwell membranes (Corning) instead of AirGels. The expansion phase and cell dissociation process were as described above. NHBE cells were loaded on Transwells at a density of ~$5\cdot10^4$ cells per well. When they reached confluence, they were transitioned to ALI culture conditions, i.e. with PneumaCult™-ALI medium (Stemcell Technologies) on the basal side and air on the apical side.

#### Live staining

To label mucus in live AirGels, we used jacalin conjugated to fluorescein (Vector Laboratories). We prepared a 50 µg/ml solution and loaded it in the lumen. We stored the chips for 30 min in a cell culture incubator before aspirating all fluid from the lumen. In addition, in order to assess epithelial integrity, we loaded a 4 kDa fluorescent dextran solution in the lumen of an 11-day-old chip and incubated it for 30 min. We then verified that all signal was localized in the lumen of the chip.

#### Immunofluorescence

All steps were performed at room temperature. First, differentiated cells in AirGels were fixed with either 4% paraformaldehyde (PFA, Electron Microscopy Sciences) or methacarn, when we wanted to better preserve extracellular mucus. Methacarn was made fresh before every use as follows: 1 part glacial acetic acid (Sigma-Aldrich), 3 parts chloroform (PanReac AppliChem), 6 parts anhydrous methanol (Sigma-Aldrich). Regardless of the chemical used, the fixation step lasted for 15 min. PFA-fixed cells were then permeabilized with a 0.2% Triton X-100 solution (VWR Life Science) for 20 min. Then, we exposed all cells (i.e. PFA- and methacarn-fixed) to a blocking solution consisting of 1% bovine serum albumin (Sigma-Aldrich) during 45 min. Afterwards, we added solutions of primary antibodies to each AirGel and incubated them for 1h. In case of PFA-fixed cells, we used rabbit anti-MUC5AC (1:100, Abcam) and mouse anti-acetylated alpha tubulin (1:250, Sigma-Aldrich); for methacarn-fixed cells, we used the same anti-MUC5AC, together with rabbit anti-MUC5B (1:100, Sigma-Aldrich). This was followed by the labeling with secondary antibodies, during 1h in the dark. More specifically, we used goat anti-rabbit IgG H&L Alexa Fluor 488 (1:200, Abcam) and goat anti-mouse IgG H&L Alexa Fluor 594 (1:200, ThermoFisher). Finally, nuclei were counterstained for 10 min with DAPI (1:1000, Sigma-Aldrich); in addition, in PFA-fixed cells, actin was stained with Phalloidin Atto 655 (1:40, Sigma-Aldrich) for 10 min.

#### Sample preparation before lightsheet imaging

To perform lightsheet microscopy on AirGels, we needed to extract the ECM gel and cells from the PDMS chip. After fixation and staining, we filled in the lumen with a 1% low-melt agarose solution in order to ensure structural integrity of the airway. We let it solidify; using tweezers, we could then carefully detach the PDMS from the glass; indeed, since the surface of the 3D printed mold was not perfectly smooth, plasma bonding was not irreversible, which we could leverage for ECM extraction. We then used a scalpel and a spatula to release the piece of ECM from the PDMS chip and later embedded it in 1% low-melt agarose. While the agarose was still liquid, we aspirated the whole gel into a 1 ml syringe (Omnifix-F, B. Braun), whose tip had previously been cut out. After the agarose solidified, we could then use the plunger to freely push the fixed AirGel in and out of the syringe, in order to image it with SPIM.

#### Microscopy

To image AirGels at low magnification (Figures 1c and d), we used a Nikon TiE epifluorescence microscope equipped with a Hamamatsu ORCA Flash 4 camera and either a 10x objective with N.A. of 0.25 or a 4x objective with N.A. of 0.1. For full channel cross-sectional imaging (Figure 1e), we used a Zeiss Lightsheet Z1 dual sided selective plane illumination microscope (SPIM). It was equipped with PCO Edge 5.5 cameras and a 5x magnification objective with N.A. of 0.16. All the other visualizations were acquired with a Nikon Eclipse Ti2-E inverted microscope coupled with a Yokogawa CSU W2 confocal spinning disk unit and equipped with a Prime 95B sCMOS camera (Photometrics). We either used a 20x water immersion objective with N.A. of 0.95, or a 40x water immersion objective with N.A. of 1.15. We used Imaris (Bitplane) for three-dimensional rendering of lightsheet z-stack pictures and Fiji for the display of all the other images[7].

#### Single-cell RNA-seq

##### Sample processing and sequencing

Three AirGels differentiated for 35 days were pooled to perform single cell RNA sequencing. The AirGels were washed three times with PBS from the apical and basal sides before carefully detaching the PDMS chip from the dish. Epithelia were removed together with their ECM from the chip using forceps and placed in dissociation buffer (300 µl Protease from Bacillus Licheniformis (100 mg/ml, Sigma), 3 µl DNase I (10 mg/ml, Roche), 30 µl EDTA (0.5 M, Sigma), 30 µl EGTA (0.5 M, BioWorld), 237 µl sterile PBS and 900 µl Accumax (Brunschwig)). Incubation was performed for 45 min at 37 °C except for centrifugation and pipetting steps which were performed at room temperature. Initially, the piece of gel and attached cells was disrupted by pipetting up and down 50 times every 5 min with a 200 µl filtered pipette tip. After the first 20 min of incubation, the cells were mostly detached from the gel and the cell suspension was centrifuged for 5 min at 400x g, after which the supernatant was removed. The residual volume (approximately 20 µl) was pipetted up and down 50 times every 5 min to disrupt cellular aggregates, this time with a 10 µl filtered pipette tip. Finally, the cell suspension of all three chips was combined and topped up to 1 ml with pre-cooled 10 % BSA (Sigma-Aldrich) in PBS. From this point, all steps were performed on ice or at 4 °C. The cells were centrifuged for 10 min at 400x g. The supernatant was carefully removed and the cells were resuspended in 0.04 % molecular grade BSA in PBS and filtered through a 40 µm Flowmi cell strainer (Bel-Art). The cell suspension was centrifuged once more at 400x g for 10 min. The supernatant was removed and the cells were resuspended in 50 µl 0.04% molecular grade BSA in PBS. The cell count was determined and the cells were immediately taken to the sequencing facility at EPFL (GECF).

HBE cells were then washed once in PBS 10% BSA and then once in PBS 0.04% BSA. After filtration through a 40 µm Flowmi strainer, cells were resuspended in PBS 0.04% BSA, checked for absence of significant doublets or aggregates, and loaded into a Chromium Single Cell Controller (10X Genomics) in a chip together with beads, master mix reagents (containing RT enzyme and poly-dt RT primers) and oil to generate single-cell-containing droplets. Single-cell Gene Expression libraries were then prepared using Chromium Single Cell 3' Library & Gel Bead Kit v3.1 (PN-1000268) following the manufacturer's instruction (protocol CG000315 Rev C). Quality control was performed with a TapeStation 4200 (Agilent) and QuBit dsDNA high sensitivity assay (Thermo) following manufacturer instructions. With this procedure, the cDNAs from distinct droplets harbor a distinct and unique 10X "cell barcode".

Sequencing libraries were processed using an Illumina HiSeq 4000 paired-end Flow Cell and sequenced using read lengths of 28 nt for read1 and 91 nt for read2, at a depth of ca 60k reads/cell.

**scRNA-seq analysis**
The Cell Ranger Single Cell Software Suite v6.1.1 was used to perform sample demultiplexing, barcode processing, and 3' gene counting using 10X Genomics custom annotation of human genome assembly GRCh38[8]. Count matrices were further processed with Seurat (version 4.1.0)[9]. All cells with less than 1,000 detected genes per cell were filtered out. Moreover, cells with more than 25% reads mapping to mitochondrial genes were removed yielding 8,651 cells passing QC. After filtering, data were default normalized and the 2,000 most variable genes identified. The expression levels of these genes were scaled before performing PCA. The following covariates were regressed out: number of UMIs and percent of mitochondrial reads UMAP dimensionality reduction was performed using the first 25 dimensions of the PCA and resolution set to 0.175. Cell subsets were identified based on transcriptional signatures previously identified by Plasschaert, Žilionis *et al* [2]. One subset was comprised of cells with a shared signature between "Ciliated" and "Secretory" cells, with a total of 640 cells, indicative of doublets and were thus removed. The remainder cells, numbering 8,011 were re-embedded as described above (resolution = 0.15). GO analysis was performed for differentially upregulated genes per cluster using *TopGO*[10].

#### Quantification of cilia beating frequency

We filled the lumens of AirGels with a 1:500 solution of yellow-green carboxylated fluorescent beads with a 2 µm diameter (FluoSpheres, Life Technologies). We incubated them for 1h; in this time interval, despite the flow generated by ciliary beating, some beads were able to settle down and attach to cilia. We then removed all fluid from the lumen and brought the chips to the spinning disk confocal microscope. We then selected small regions of interest (50 x 50 pixels, i.e. 27.5 µm x 27.5 µm) around individual beads and recorded videos at 100 frames per second. Using Matlab R2016b (Mathworks), we computed the mean intensity of each frame over time. We computed the fast Fourier transform of each mean intensity signal, which we then used to obtain single-sided power spectra. We only kept frequencies between 1 and 30 Hz, thereby getting rid of artifacts. We finally looked for the frequency with maximal amplitude in the power spectrum, which corresponded to the cilia beating frequency.

#### Quantification of mucociliary clearance

Like for CBF quantification, we loaded a 1:500 solution of 2 µm FluoSpheres in the lumen of AirGels. We immediately visualized them with the spinning disk confocal microscope. We recorded 10 s videos at a rate of 10 frames per second. Then, tracked the trajectory of individual beads with the Fiji plugin TrackMate[11], using the built-in simple LAP tracker. We wrote a script in a Jupyter Notebook to compute the velocity (track displacement over track duration) of each particle[12].

#### Bacterial strains, plasmids and culture conditions

We used *Pseudomonas aeruginosa* PAO1 (WT or mutants, listed in Supplementary Table 1) for all the infection experiments. Most strains were made to constitutively express the fluorescent protein mScarlet following a published protocol using the plasmids listed in Supplementary Table 2[13]. The backbone plasmid pUC18t-Mini Tn7 with gentamycin resistance was purchased from Addgene and isolated from *E. coli* XL10 Gold by GeneJET Plasmid Miniprep Kit (Thermo Fisher). The isolated plasmid was digested with the restriction enzymes HindIII and BamHI. The Ptet promoter was amplified by PCR using *P. aeruginosa* PAO1 genomic DNA and the mScarlet gene was amplified from a pre-existing plasmid. The Ptet promoter and mScarlet was then fused via Fusion PCR by overlapping extension. The resulting extended product was digested with HindIII and BamHI, then ligated to the digested pUC18t-MiniTn7 Gm backbone. Since this plasmid included a gentamycin resistance cassette, we grew the fluorescent PAO1 strains overnight in LB medium with 30 µg/ml gentamycin. The next morning, we diluted the stationary cultures 1:1000 in plain LB and let them grow 3-4h before infecting AirGels.

**Supplementary Table 1: strains used in this study**

| Strain | Relevant characteristics | Source / Reference |
|---|---|---|
| *Pseudomonas aeruginosa* PAO1 (ATCC 15692) | WT PAO1 | [14] |
| PAO1 mScarlet | WT PAO1 with constitutive chromosomal mScarlet expression | This study |
| PAO1 *Δxcp* | *xcpP* to *xcpZ* chromosomal deletion (called DZQ40 in the original study) | [15] |
| PAO1 *Δxcp* mScarlet | PAO1 *Δxcp* with constitutive chromosomal mScarlet expression | This study |
| PAO1 *ΔfliC* | In-frame deletion of PA1092 | [16] |

| | | |
|---|---|---|
| PAO1 *ΔfliC* mScarlet | PAO1 *ΔfliC* with constitutive chromosomal mScarlet expression | This study |
| PAO1 *ΔpilA* | In-frame deletion of *pilA* | [17] |
| PAO1 *ΔpilA* mScarlet | PAO1 *ΔpilA* with constitutive chromosomal mScarlet expression | This study |
| PAO1 *ΔpilT* | *pilT*::Tn5 | [18] |
| PAO1 *ΔpilT* mScarlet | PAO1 *ΔpilT* with constitutive chromosomal mScarlet expression | This study |
| PAO1 *ΔpilH* | In-frame deletion of PA0409 | [19] |
| PAO1 *ΔpilH* mScarlet | PAO1 *ΔpilH* with constitutive chromosomal mScarlet expression | This study |
| PAO1 *ΔfliC ΔpilH* | In-frame deletion of PA1092 and PA0409 | [20] |

**Supplementary Table 2: Plasmids used in this study**

| **Plasmid** | **Source** | **Reference** |
|---|---|---|
| pTNS2 | Addgene 64968 | [21] |
| pUC18T-mini-Tn7T-Gm-Ptet_mScarlet | Addgene 63121 with Ptet promoter fused to mScarlet | This study |

#### Infection of AirGels

The night before infection, AirGels were stained with the plasma membrane dye CellMask Deep Red (Life Technologies). Aside from the infection assay shown in Figure 3a and Supplementary Video 2, which was performed in a CF AirGels, all infections were run with cells from healthy donors. The dye was diluted to 5 µg/ml and loaded in both the apical and basal compartments. The next morning, the lumen was again exposed to air for 3-4h. Mucus was stained with jacalin as described above, and all luminal fluid was then aspirated. Finally, we infected AirGels with mScarlet *P. aeruginosa*. We measured the optical density of our exponential bacterial cultures and centrifuged them for 2-3 min at 5000 rpm. We discarded the supernatant and resuspended the pellet in D-PBS to reach an optical density value of approximately 3. We then loaded 0.5 µl of bacterial culture in the lumen of AirGels (this small volume allowed for ALI maintenance). The resulting multiplicity of infection was approximately 10. For the infection shown in Figure 3a, we started with a stationary *P. aeruginosa* culture that we diluted in D-PBS to an optical density of ~0.035. We then dipped a sterile toothpick in the culture and lightly touched the inlet of an AirGel with it in order to deposit bacteria while maintaining the ALI. This second method may mechanically compromise the epithelium with the toothpick, we therefore opted for the first one in most experiments.

The chips were then placed in an UNO-T-H-CO2 stage-top incubator (Okolab) for temperature, humidity and $CO_2$ control. The environment was maintained at 37°C and 5% $CO_2$, and connected to a bottle of Milli-Q water for humidification. Since condensation frequently appears on the PDMS chip during imaging, we placed pieces of Kimtech Science™ Kimwipes™ (Kimberly-Clark Professional) in the inlet ports of AirGels; this prevented dripping water from disrupting the ALI conditions. We visualized the infection progress over time with the aforementioned spinning disk confocal microscope. For WT, *ΔpilT* and *ΔpilH*, we repeated the infections to reach *N* = 3 replicates per condition. The AirGels for all 3 replicates were all made from the same healthy donor and were between 33- and 38-day-old at the time of infection.

#### Colonization of extracted mucus

We isolated mucus from 8.5-month-old NHBE cells grown on 0.4 µm pore size polyester Transwell membranes (Corning). To do so, we immersed the apical side of the membranes in a jacalin-fluorescein solution (50 µg/ml in D-PBS) and we placed them in a cell culture incubator for 30 min. We then collected all fluid from the apical side with a pipette and dispensed 12.5 µl into 4 mm PDMS gaskets bonded to a glass-bottom dish (1.5 coverslip, glass diameter 20 mm, MatTek). We filled the space around the PDMS gasket with D-PBS to prevent dehydration of the mucus. We then centrifuged an exponential *P. aeruginosa* mScarlet culture at 5000 rpm for 3 min and resuspended them in D-PBS before loading 15 µl on the labeled mucus. We then placed the dish in the stage-top incubator and imaged the bacteria and mucus with the confocal spinning disk microscope described above.

#### Twitching motility on mucus

We infected a 4-month-old HBE Transwell with *P. aeruginosa* mScarlet as follows. We loaded 3.3 µl of early exponential culture (~$10^5$ colony-forming units) on the apical side of the HBE culture, which had been labeled with fluorescent jacalin. We carefully took the Transwell insert out of the cell culture plate using sterile tweezers and we placed it on a glass-bottom dish (1.5 coverslip, MatTek). We then placed the dish in a stage-top incubator and recorded time-lapse videos of twitching bacteria with our spinning-disk confocal microscope. Because of the lack of culture medium in the visualization setup, recordings could not last long and would dehydrate within minutes.

#### Biofilm image acquisition and analysis

We acquired z-stack of infected AirGels over a 35 µm-deep range at different time points ($t$ = 0h, 2h, 4h and 5.5h ± 0.5h). All the image analysis steps were done in Jupyter Notebooks[12]. To estimate the growth rates of different strains, we summed the total fluorescence intensity over all slices and normalized the obtained value with respect to time $t$ = 0h.

Since the AirGel surface is curved, for all subsequent steps, we projected images in 2D using the maximal intensity projection tool in Fiji in order to facilitate downstream analysis. We started by quantifying the sizes of bacterial clusters. First, we visually inspected the pictures: if there were large intensity variations (e.g. in case of a mix of dim single cells and bright clusters), we saturated bright pixels to 1.5 times the mean intensity of the picture. We then segmented the pictures using Otsu thresholding (from the 'opencv' Python package[22], version 4.5.4.60) and visually assessed the result. In the rare cases where the segmentation was not deemed satisfactory (i.e. if some features were not detected properly or if there was too much noise), a simple threshold was manually selected. The pictures were then closed and filtered; more specifically, we removed any object smaller than ~6 µm$^2$ (20 pixels), which approximately corresponds to the area of a single cell. We then obtained the area of each cluster using the function 'regionprops' ('scikit-mage' Python package[23], version 0.19.2), which calculates properties of segmented objects in binary pictures. We calculated the mean cluster area for each replicate; then, for each condition, we plotted the maximum, minimum and mean of the means (e.g. Figure 3b). We also computed and plotted the proportion of aggregates larger than 100 µm$^2$ (Supplementary Figure 2a).

We then quantified colocalization between mucus and bacteria. The segmentation and filtration of mucus pictures was identical as for bacterial clusters. Then, using the logical '&' function, we identified the pixels that were common between the binary pictures from the bacterial and mucus channels. With 'regionprops', we obtained the areas of these common zones and we normalized them to the total area of mucus. Thus, we could find the proportion of mucus that was covered in bacteria. We finally calculated the proportion of mucus devoid of bacteria as follows: 1 – (proportion of mucus covered in bacteria).

To quantify the contraction of a patch of mucus, we first canceled the effects of drift by registering the images in Fiji using the 'Correct 3D drift' plugin. We then manually tracked the displacement of $N = 7$ reference features with the Fiji plugin 'Manual Tracking'. We loaded the trajectories in a Jupyter Notebook and calculated the distances between each pair of positions over time. We finally normalized the resulting data to the initial distances and plotted them, along with the mean and standard deviation at each time point (Figure 4d).

Finally, we also measured mucus shrinkage over time for WT, *ΔpilH* and an uninfected control AirGel. To do so, we used images from 30 min timelapses (the starting point of the timelapses differed: 6 h 10 min for WT, 2 h 30 min for *ΔpilH*, and 8 h 5 min for the negative control). We segmented and quantified mucus areas as described above for each time point, and normalized it to the initial area (Figure 6c & Supplementary Video 7).

#### iSCAT-based quantification of type IV pili retraction frequency

*P. aeruginosa* were grown as previously described[20]. Briefly, an overnight culture was obtained from a single colony and grown in LB at 37°C with 290 rpm shaking. The overnight culture was diluted 1:500 or 1:1000 and grown for 2 to 3 hours to obtain a mid-exponential phase culture. For surface-grown cells, 100 µl of the mid-exponential phase cell suspension were plated on LB 1 % agarose, grown for 3 h at 37 °C and harvested in 500 µl LB by gentle scraping. Cells were diluted to OD600 0.02 to 0.05 prior to visualization. Both liquid- and solid-grown cells were either loaded on 500 µm x 140 µm PDMS microchannels or in 6 mm PDMS gaskets. Cells sticking to the surface were visualized without flow with iSCAT and movies were recorded at 10 fps for either 2 min, 1 min or 30 s. Raw iSCAT images were processed as described previously[3,20]. Individual movies were manually analyzed using Fiji[7] by counting the total number of TFP in each cell as well as the number of TFP retractions represented by tensed TFP. The residence time of each cell on the coverslip was also recorded. For each cell we computed the retraction frequency by dividing the total number of retractions by the residence time of the cell on the coverslip. Finally, we computed a bootstrap median retraction frequency and 95% confidence interval by pooling the data obtained by all three biological replicates. Data analysis was performed using Matlab R2020a (Mathworks).

#### Statistical analysis

All statistical tests were run in Python using Jupyter Notebooks[12]. Independent or paired-samples Student t-tests were performed with Bonferroni correction using the function 'add_stat_annotation' from the statannot package[24] (version 0.2.3). One-way ANOVAs were run using the function 'f_oneway' in the 'stats' module from SciPy[25] (version 1.7.3). When the ANOVA result rejected the null hypothesis, we followed up with a post-hoc Tukey test using 'stats.multicomp.pairwise_tukeyhsd' from the 'statsmodels' module[26].

#### Computational model of mucus remodeling by T4P

We refer the reader to our previous work[27,28] for the general theory on the kinematics of the surface and the volume of a 3D soft body, focusing on cell-laden microtissues. Specific considerations on the implementation of this work are introduced in the following formulations.

##### 1. Kinematics

Let $V$ be a fixed reference configuration of a continuum body $\mathcal{B}$. We use the notation $\chi: V \rightarrow \mathbb{R}^3$ for the deformation of body $\mathcal{B}$. A motion $\chi$ is the vector field of the mapping $\boldsymbol{x} = \chi(\mathbf{X})$, of a material point in the reference configuration $\mathbf{X} \in V$ to a position in the deformed configuration $\boldsymbol{x} \in v$. The kinematics of a material point are described by

$$\mathbf{u}(\mathbf{X}, t) = \boldsymbol{x}(\mathbf{X}, t) - \mathbf{X} \quad \text{(S1)}$$

where $\mathbf{u}(\mathbf{X}, t)$ is the displacement vector field in the spatial description. The kinematics of an infinitesimal bulk element are described by

$$\mathbf{F}(\mathbf{X}, t) = \frac{\partial \chi(\mathbf{X}, t)}{\partial \mathbf{X}} = \nabla_{\mathbf{X}} \boldsymbol{x}(\mathbf{X}, t) \quad \text{(S2)}$$

$$\mathbf{F}^{-1}(\boldsymbol{x}, t) = \frac{\partial \chi^{-1}(\boldsymbol{x}, t)}{\partial \boldsymbol{x}} = \nabla_{\boldsymbol{x}} \mathbf{X}(\boldsymbol{x}, t) \quad \text{(S3)}$$

where $\mathbf{F}(\mathbf{X}, t)$ and $\mathbf{F}^{-1}(\boldsymbol{x}, t)$ are the deformation gradient and inverse deformation gradient, respectively. Note that $J(\mathbf{X}, t) = \mathrm{d}v/\mathrm{d}V = \det \mathbf{F}(\mathbf{X}, t)$ is the Jacobian determinant defining the ratio of a volume element between material and spatial configuration.

A motion of an arbitrary differential vector element can be mapped by the deformation gradient $\mathbf{F}$. However, a unit normal vector $\mathbf{N}$ in the reference configuration cannot be transformed into a unit normal vector $\mathbf{n}$ in the current configuration via the deformation gradient[29], motivating us to develop the kinematics of an infinitesimal surface element[30]. Note that we utilize $\{\widehat{\bullet}\}$ to denote the surface quantity bounded by outer surface denoted as $\partial\Omega_0$.

$$\widehat{\mathbf{F}}(\mathbf{X}, t) = \frac{\partial \chi(\mathbf{X}, t)}{\partial \mathbf{X}} \cdot \widehat{\mathbf{I}} = \widehat{\nabla}_{\mathbf{X}} \boldsymbol{x}(\mathbf{X}, t) \quad \text{(S5)}$$

$$\widehat{\mathbf{F}}^{-1}(\boldsymbol{x}, t) = \frac{\partial \chi^{-1}(\boldsymbol{x}, t)}{\partial \boldsymbol{x}} \cdot \hat{\boldsymbol{\imath}} = \widehat{\nabla}_{\boldsymbol{x}} \mathbf{X}(\boldsymbol{x}, t) \quad \text{(S6)}$$

where $\widehat{\mathbf{F}}(\mathbf{X}, t)$ and $\widehat{\mathbf{F}}^{-1}(\boldsymbol{x}, t)$ are the deformation gradient and inverse deformation gradient, respectively. Note that $\widehat{\mathbf{I}} = \mathbf{I} - \mathbf{N} \otimes \mathbf{N}$ and $\hat{\boldsymbol{\imath}} = \boldsymbol{i} - \mathbf{n} \otimes \mathbf{n}$ are the mixed surface unit tensors, where $\mathbf{I}$ and $\boldsymbol{i}$ are the unit tensors, and $\mathbf{N}$ and $\mathbf{n}$ are the outward unit normal vectors, in reference and current configuration, respectively. Note that $\hat{J}(\mathbf{X}, t) = \mathrm{d}a/\mathrm{d}A = |\mathrm{cof}\, \mathbf{F} \cdot \mathbf{N}|$ is the Jacobian determinant defining the ratio of an area element between material and spatial configuration.

#### 2. Equilibrium

The total potential energy functional $W(\chi)$ is defined as:

$$W(\chi) = \int_{\Omega_0} \Psi(\mathbf{F}, \chi; \mathbf{X}) \mathrm{d}V + \int_{\partial\Omega_0} \widehat{\Psi}(\widehat{\mathbf{F}}, \chi; \mathbf{X}) \mathrm{d}S - \int_{\Omega_0} \mathbf{B} \cdot \mathbf{u}(\chi; \mathbf{X}) \mathrm{d}V - \int_{\partial\Omega_0} \mathbf{T} \cdot \mathbf{u}(\chi; \mathbf{X}) \mathrm{d}S \quad \text{(S9)}$$

where $\Psi$ and $\widehat{\Psi}$ are strain energies in bulk and on surface, and $\mathbf{B}$ is the reference body force and $\mathbf{T}$ is the surface traction. An equilibrated configuration is obtained by minimizing this functional considering all admissible deformations $\delta\chi$. It is important to note that the strain energies $(\Psi, \widehat{\Psi})$ can be varied depending on the bacterium and mucus models so that the following sections can be written in a single formulation for brevity, and the specific strain energies are to be defined in section 4.

Following the derivation presented in [31], we can finally arrive at a set of localized force balance equations. Neglecting the inertial effect, the local form of linear and angular momentum balances for bulk and surface are defined by

$$\nabla_{\mathbf{X}} \cdot \mathbf{P} + \mathbf{B} = \mathbf{0} \quad \text{in } V \quad \text{(S10)}$$

$$\hat{\nabla}_{\mathbf{X}} \cdot \hat{\mathbf{P}} + \mathbf{T} - \mathbf{P}\mathbf{N} = \mathbf{0} \quad \text{on } S \tag{S11}$$

$$\mathbf{u} = \breve{\mathbf{u}} \quad \text{on } S_u \tag{S12}$$

$$[\![\hat{\mathbf{P}}\hat{\mathbf{N}}]\!] = 0 \quad \text{on } L$$

where $\breve{\mathbf{u}}$ is the prescribed displacement on the boundary $S_u$, $\hat{\mathbf{N}}$ is the bi-normal vector to the boundary curve, and $[\![\bullet]\!]$ indicates summation over surfaces intersecting on boundary curves[30].

#### 3. Weak Form

For the finite element implementation, we need to obtain the weak form for our problem. By adding the constraint that the first variation of the total potential energy must be equal to zero $\delta W(\chi) = 0$, we obtain a weak form statement as

$$G = \int_{\Omega_0} \mathbf{P} : \nabla_{\mathbf{X}} \delta\mathbf{u} \, \mathrm{d}V + \int_{\partial\Omega_0} \hat{\mathbf{P}} : \hat{\nabla}_{\mathbf{X}} \delta\mathbf{u} \, \mathrm{d}S - \int_{\Omega_0} \mathbf{B} \cdot \delta\mathbf{u} \, \mathrm{d}V - \int_{\partial\Omega_0} \mathbf{T} \cdot \delta\mathbf{u} \, \mathrm{d}S = 0 \quad \forall \, \delta\mathbf{u} \tag{S13}$$

where $\delta\mathbf{u}$ is the admissible deformation field.

We employed the open-source platform FEniCS[32], to implement the finite element simulation. We used the Scalable Nonlinear Equations Solvers (SNES) from the open-source toolkit PETSc[33], which provides numerical computations of a Newton-type iterative procedure to solve the nonlinear variational problem. Note that the value of $\gamma$ should be ramped from zero to its prescribed value for numerical stability as the problem is highly nonlinear.

#### 4. Constitutive Relations

To relate the active stress with deformation, we must specialize our choice for the strain energies in the bulk and on the surface. For the deformation of compressible tissue, we consider a passive bulk energy $\Psi^p$ that captures the permanent elasticity of the collagen network, and for the contribution of bacterium contractility, we can consider the active surface energy $\hat{\Psi}^a$ that accounts for the action of the bacterium on the surface of mucus tissue.

##### 4.1. Passive mucus model

The passive strain energy density $\Psi^p$ describes the elasticity of mucus tissue. We consider the mucus as a soft, highly deformable and highly compressible hyperelastic material, but we neglect it’s biphasic and viscoelastic nature in terms of energy dissipation. We choose the compressible Neo-Hookean model [28,29] for the mucus.

$$\Psi^p = \frac{K}{2}(J-1)^2 + \frac{G}{2}(I_1 - 3 - 2\ln J) \tag{S14}$$

where $K$ and $G$ are the bulk and shear moduli.

##### 4.2. Active model for the contractile action of bacteria

We assume that the bacteria-mucus interaction can be described through a surface strain energy generating constant surface stresses similar to fluid-like surface tension[27,28]. Bacteria exert a contractile force on the periphery of the mucus, and we recapitulate this action through an active surface energy $\hat{\Psi}^a$. We postulate that the surface energy $\hat{\Psi}^a$ is a function of the change of the surface area $\hat{J}$.

$$\hat{\Psi}^a = \gamma\hat{J} \quad \text{(S15)}$$

where $\gamma$ is a surface contractile modulus (energy per unit area) representing the contractility of bacterium on the surface at the equilibrium state. It is important note that we consider no bulk contractility due to the bacteria as results verify their presence only on the periphery of the mucus.

##### 4.3. Energy penalization

As at high level of contraction the bacteria are bound to jam, barring the additional contraction of the mucus (even if the material itself can accommodate it), we have to enforce this jamming transition. Assuming that we know the initial surface concentration of bacteria we enforce the kinematic constraint via energy penalization. From the experimental observation, we enforce the surface area of deformed mucus tissue cannot be smaller than a ratio ($\hat{J}_{pen}$) of the initial surface area. An appropriate energy penalization $\hat{\Psi}_{pen}$ for enforcing the prescribed surface condition is given by

$$\hat{\Psi}_{pen} = \frac{P}{2}\langle \hat{J}_{pen} - \hat{J}\rangle^2 \quad \text{(S16)}$$

where $P$ is the penalty parameter (energy per unit area), and $\langle \bullet \rangle$ is the Macaulay brackets that used to describe the ramp function,

$$\langle x \rangle = \begin{cases} x & (x > 0) \\ 0 & (x \leq 0) \end{cases} \quad \text{(S17)}$$

#### 5. Finite Element Simulation

The reference (undeformed) state corresponds to a state where the active contractile moduli are set to zero. Experimentally, this reference state corresponds to the initial state of the mucus right after the mixing of mucus and bacterium and before the application of forces by encapsulated bacterium. The reference configuration for the finite element simulations represents the geometry shown in Figure 5b. The entire surface is allowed to actively contract through increasing the surface contractile modulus up to an equilibrium value. The final (deformed) state is defined when the surface contractile moduli $\gamma$ reaches its prescribed value, and no external loads are applied. Experimentally, this corresponds to the equilibrium state of the mucus. The final configurations represent the equilibrium states.

#### 6. Parameter Calibration

The parameters of the model are the bulk and shear moduli, $K$ and $G$, and surface contractile modulus, $\gamma$, penalty parameter, $P$ and penalty surface ratio, $\hat{J}_{pen}$. There is a unique relationship between $K$, $G$, and Poisson's ratio $\nu$, allowing to interchangeably use $\nu$ in place of $K$ for the calibration procedure. We selected a set of parameters: $G = 1.0$ Pa, $\nu = 0.1$, $\gamma = 0.03$ nN/µm and $P = 1.0$ nN/µm. The corresponding bulk and elastic moduli were $K = 0.9$ Pa and $E = 2.2$ Pa within the range of reported values for mucus[34,35].
